# CellART: a unified framework for extracting single-cell information from high-resolution spatial transcriptomics

**DOI:** 10.64898/2026.09.03.749294

**Authors:** Yuheng Chen, Yuyao Liu, Zhiwei Wang, Yeqin Zeng, Zitong Chao, Peiqi Jiang, Hao Chen, Jiguang Wang, Jiashun Xiao, Can Yang

## Abstract

Understanding how different cell types assemble into tissues and organs, as well as how they interact to transmit and receive biological signals, is essential for advancing biomedical and biological research. Recent advancements in spatial transcriptomics (ST) technologies have opened new avenues for investigating biological systems by achieving subcellular spatial resolution. Since cells are the fundamental units of life, extracting single-cell information from high-resolution ST data is crucial. However, existing ST platforms often capture sparse transcript counts per spot or measure only a limited number of genes, complicating the extraction of comprehensive single-cell information. In this study, we introduce CellART, a unified framework designed to extract single-cell information across diverse high-resolution ST platforms, including VisiumHD, Xenium, MERFISH, and Stereo-seq. By leveraging multimodal data, such as staining images, spatial transcriptomics data, and single-cell RNA sequencing references, CellART simultaneously performs cell segmentation and cell type annotation through a seamless integration of deep learning and probabilistic modeling. We demonstrate the efficiency, generalizability, and robustness of CellART across various high-resolution spatial transcriptomics platforms, capable of processing datasets containing millions of spots. Comprehensive experiments validate the biological relevance and accuracy of the recovered cellular information within spatial configurations. Notably, we highlight the utility of CellART in breast and colorectal cancer datasets, showcasing its ability to fully leverage high-resolution ST data. By enhancing cellular resolution, CellART facilitates the identification of transient cancer cell states and immune cell subtypes. Furthermore, CellART enables investigations into cancer-immune cell communication, uncovering both established interactions and novel ligand-receptor pairs. The outputs of CellART are compatible with widely used community tools, facilitating a variety of downstream analyses.

## Introduction

A central question in biological and medical research is how cells are organized in space and how they interact. Spatial transcriptomics (ST) has transformed this field by enabling the measurement of gene expression profiles within their natural spatial context [1, 2]. Recent advancements in ST technologies have significantly improved the spatial resolution of ST data to the subcellular level. Sequencing-based ST approaches, such as VisiumHD [3], Seq-Scope [4], Open-ST [5], and Stereo-seq [6], achieve spatial resolutions ranging from 0.5 to 2 *µ*m, while imaging-based techniques like 10x Xenium [7], MERFISH [8], and CosMx [9] offer resolutions as fine as 0.1 to 0.2 *µ*m. Given that cells are the fundamental units of living organisms, it is essential to extract information at the single-cell level from the subcellular measurements provided by these high-resolution ST technologies. This enables a deeper characterization of cellular organization and communication, ultimately enhancing our understanding of biological processes and disease mechanisms [10, 11, 12, 13].

However, recovering single-cell information from high-resolution ST data presents significant challenges due to imperfect subcellular measurements. Sequencing-based ST technologies often suffer from low transcriptional abundance per spot. The standard processing pipeline typically aggregates neighboring spots (e.g., combining 2 × 2 *µ*m grids into 8 × 8 *µ*m spots for VisiumHD data) and then applies deconvolution methods, such as RCTD [14] and Cell2location [15], to estimate cell type proportions within these coarsely binned spots. This pipeline cannot fully harness the power of high-resolution data, as binned spots do not accurately reflect true cell boundaries and introduce unwanted cell type mixtures in downstream analyses. While several methods, such as TopACT [16], SCS [17], and Bin2Cell [18], have been developed for analyzing high-resolution sequencing-based ST data, they fall short in recovering single-cell information due to algorithmic instability, oversimplified model assumptions, or low computational efficiency. For imaging-based ST technologies, existing approaches for single-cell recovery often exhibit limited performance. Methods such as Cellpose [19], StarDist [20], ProSeg [21], and Baysor [22] frequently yield inaccurate segmentation results owing to their reliance on single-modality inputs, whereas more recent methods like Bering [23] and Cellotype [24] require labor-intensive manual annotation, thereby constraining their scalability and broader applicability.

To address the aforementioned challenges, we present CellART, a unified framework designed to extract single-cell informAtion from high-Resolution ST data. CellART integrates information from staining images, ST samples, and single-cell RNA sequencing (scRNA-seq) references to simultaneously perform cell segmentation and cell type annotation. The core innovation of CellART lies in its approach to jointly model these two tasks through a shared high-resolution latent representation learned within a probabilistic framework, rather than treating them as independent or sequential steps. This integrated formulation allows each task to enhance the other: cell type information from the scRNA-seq reference is embedded into the shared latent space during representation learning, providing biologically informed features for segmentation. The resulting refined cell boundaries, in turn, improve expression aggregation for more accurate annotation. With its innovative model design, CellART efficiently extracts single-cell information from high-resolution ST data comprising millions of cells, generating outputs that include cell and nuclei boundaries, gene expression profiles, and cell type labels for individual cells. Its strength lies in its unified framework, which combines deep neural networks with a Poisson-based probabilistic model. This enables CellART to leverage multimodal information from high-resolution imaging, transcript measurements, and scRNA-seq reference data while scaling to large ST datasets. In this framework, CellART achieves cell segmentation and cell type annotation through three steps: learning high-resolution representations of each subcellular spot, segmenting cells through adaptive labeling of positive and negative samples, and optimizing cell type annotations using a probabilistic model to account for uncertainty. In contrast to existing approaches, CellART effectively integrates multimodal information from high-resolution ST data and scRNA-seq data through likelihood-based optimization, maximizing the use of available data and eliminating the need for extensive manual annotation.

We demonstrate the superiority of CellART across a wide range of datasets comprising different species, tissues, diseases, and high-resolution ST platforms (including both sequencing-based and imaging-based platforms). Benchmarking on both imaging-based and sequencing-based ST datasets reveals CellART’s superior accuracy and computational efficiency compared to existing methods. We further apply CellART to mouse brain coronal datasets from four distinct high resolution ST platforms: Xenium, VisiumHD, MERFISH, and Stereo-seq. CellART accurately recovers cell boundaries and cell type labels within different tissue structures, such as the hippocampus and cortical brain layers, and achieves consistent alignment of these structures across platforms. This underscores its robustness and versatility across diverse technologies. In human breast cancer, the precise cellular information extracted by CellART enables the identification of transient cancer cells, providing insights into the progression of disease states. By leveraging subcellular structures revealed by CellART to estimate RNA velocity, we observe that two ductal carcinoma in situ (DCIS) subtypes exhibit distinct pathways, with one subtype showing a greater tendency to progress into invasive tumors. CellART also demonstrated its ability to enhance resolution and reduce transcript misallocation in VisiumHD colorectal cancer datasets. By directly extracting single-cell level information from raw subcellular spots, CellART achieves a more detailed and biologically accurate reconstruction of cellular spatial organization compared to conventional binned-spot-based analyses. This capability enables downstream applications, such as the detection of immune cell subtypes and the identification of cellular niches. Building on its single-cell resolution, CellART reveals tumor-associated macrophages (TAMs) in colorectal cancer, characterizing their spatial distribution and highlighting the expression of enhanced immune-related genes. Ligand-receptor analysis further uncovers cell-cell communication between TAMs and cancer cells, providing deeper insights into the tumor microenvironment and potential mechanisms of tumor-immune interaction. We anticipate that CellART will serve as a scalable and versatile tool for analyzing high-resolution ST data.

## Results

### Overview of CellART

CellART is a statistical and computational framework designed to address the challenges of extracting single-cell information from high-resolution ST data. By employing deep learning and probabilistic modeling to effectively integrate the power of multimodal data, including spatial transcriptomics, staining images, and scRNA-seq references, it achieves a unified and trustworthy framework for cell segmentation and cell type annotation. In contrast to conventional “segment-then-annotate” pipelines that decouple cell boundary determination from cell type inference, CellART couples these tasks through a shared latent representation that encodes both spatial and transcriptomic information. This unified design ensures that representation learning, segmentation, and annotation are mutually reinforcing: the probabilistic model embeds cell type knowledge into the shared representation, the representation provides biologically informed features for segmentation, and the refined cell boundaries improve cell-level aggregation for annotation. CellART effectively bridges the gap between noisy subcellular measurements and biologically meaningful cell-level information, enabling accurate and comprehensive characterization of cellular organization across diverse ST platforms.

The CellART framework employs a three-step strategy to extract single-cell level information from high-resolution ST data (Fig. 1**a**, Supplementary Fig. S2, Methods). It begins with learning high-resolution representations for subcellular spots, which serve as the foundation for subsequent cell segmentation and cell type annotation. This step combines probabilistic modeling with a feature pyramid network (FPN) to address the noisy and sparse nature of subcellular measurements in high-resolution ST data and to capture multiscale spatial features. Moreover, by coarsely aggregating subcellular spot representations into cell-level representations based on nuclei segmentation and modeling their correspondence with cell type information from the scRNA-seq reference, CellART guides the learning of biologically meaningful representations.

**Figure 1.**
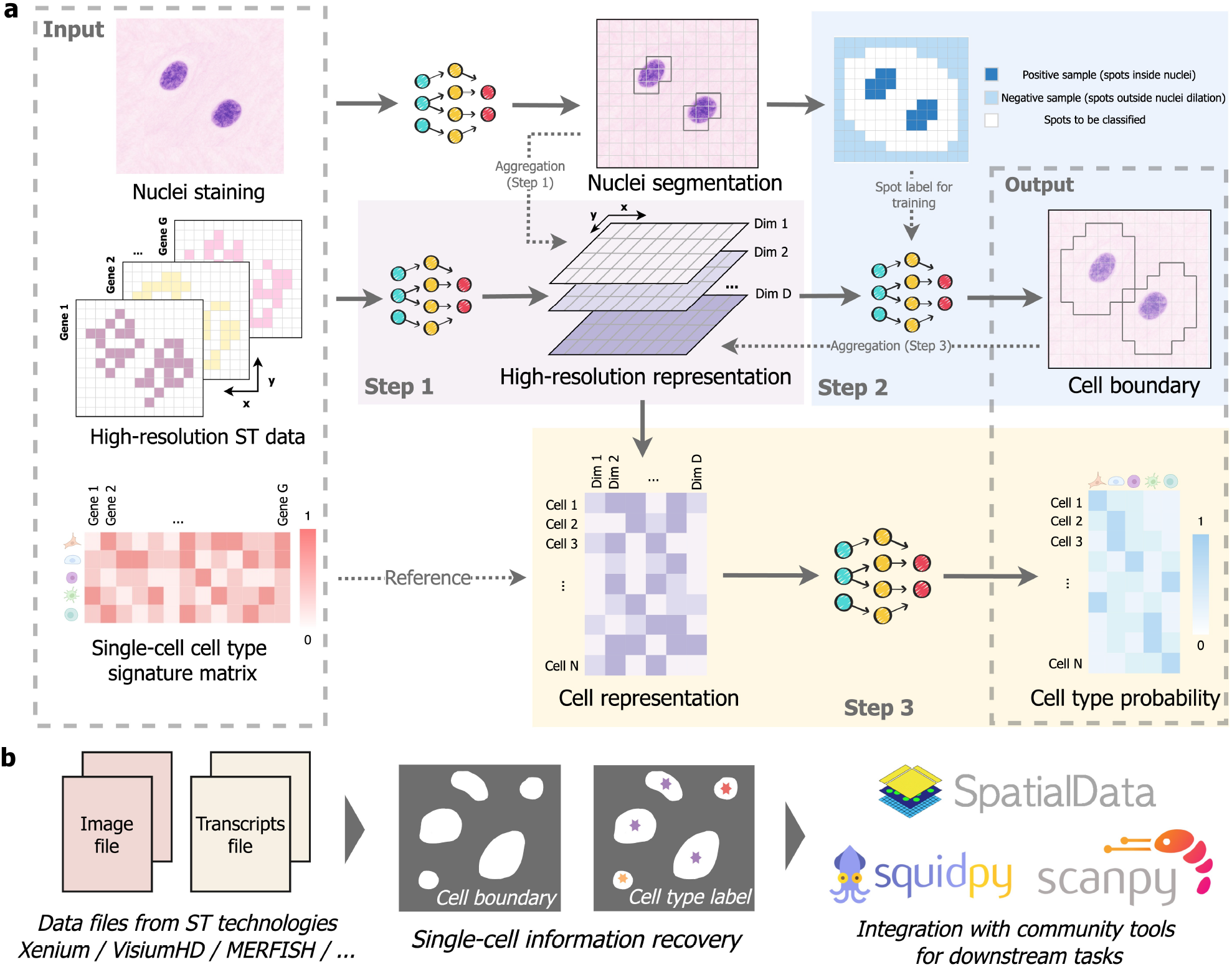
CellART framework. **a.** CellART utilizes high-resolution ST data and scRNA-seq reference data as input, producing the boundaries, expression profiles, and cell types of individual cells as output. CellART comprises three steps. Step 1: high-resolution representation learning. CellART initiates the process by learning high-resolution representations of subcellular spots through the integration of a deep neural network and a probabilistic model. Specifically, it employs a feature pyramid network (FPN) to extract detailed representations from spatial transcriptomics (ST) data, enabling the characterization of subcellular spots by incorporating information from both the spots themselves and their neighboring areas. To address the noise inherent in subcellular measurements, a probabilistic model is developed to aggregate these representations into cell-level data, linking them to scRNA-seq reference data. This likelihood-based optimization significantly enhances the quality of representation learning. Step 2: Cell segmentation. Positive samples are defined as spots within the nucleus, while negative samples are those located far from nuclei. A neural network is trained using the high-resolution representations as input and the positive and negative labels as output to classify spots, allowing for the determination of cell boundaries. Step 3: Cell type annotation. Following cell segmentation, representations and expression profiles are aggregated at the whole cell level. The probabilistic model is further optimized for accurate cell type annotation. **b**. CellART enables the extraction of single-cell level information, including cell boundaries and cell type labels, from diverse ST platforms. It is compatible with community tools, such as SpatialData, Scanpy, and Squidpy, facilitating downstream analyses.

After obtaining the representations for each subcellular spot, CellART proceeds to delineate cell boundaries by introducing a segmentation network. A key challenge in this step is the lack of ground truth labels for spots, complicating the training of the segmentation network. To address this issue, we use nuclei segmentation masks to adaptively generate positive samples (spots within nuclei) and negative samples (spots outside dilated nuclear regions) without human intervention. This approach ensures a robust distinction between cells and background. The trained segmentation network outputs the probability of each spot belonging to each cell, enabling CellART to achieve smooth and accurate cell boundary delineation.

With the cell segmentation result, CellART obtains more accurate and less noisy cell-level representations based on the cellular mask rather than the nuclear mask. It then further optimizes the initial model and employs the refined probabilistic framework to achieve precise cell type annotation. Details are included in the Methods.

CellART is broadly applicable to high-resolution ST data, including Xenium [7], VisiumHD [3], MERFISH [8], and Stereo-seq [6]. It produces user-friendly outputs optimized for downstream analyses and is fully compatible with widely used bioinformatics tools, such as SpatialData [25], Scanpy [26], and Squidpy [27] (Fig. 1**b**). For example, the cellular information inferred by CellART can be directly applied to investigate spatial gene expression patterns, infer dynamic cell states, and identify cell niches, providing deeper insights into tissue organization and function. This compatibility with existing workflows makes CellART a practically useful tool for analysis of high-resolution ST datasets. The CellART software is publicly available at https://github.com/YangLabHKUST/CellART.

The Results below are organized in two parts. We first establish CellART’s methodological performance through systematic benchmarking and cross-platform validation on mouse brain datasets from four ST platforms, demonstrating its accuracy, robustness, and computational efficiency. We then apply CellART to breast and colorectal cancer datasets to illustrate the biological insights that its unified framework and enhanced single-cell resolution can enable in disease contexts.

### Benchmarking studies highlight the superior performance of CellART in recovering single-cell information

To benchmark CellART’s performance, we conducted a series of comparative analyses using both imaging-based and sequencing-based high-resolution ST datasets (Fig. 2). The benchmarking was structured in three parts: first, evaluating segmentation performance on representative imaging-based (Xenium) and sequencing-based (VisiumHD) platforms; second, benchmarking annotation accuracy using a paired dataset through consistency metrics; and finally, assessing computational efficiency to validate scalability across large datasets.

**Figure 2.**
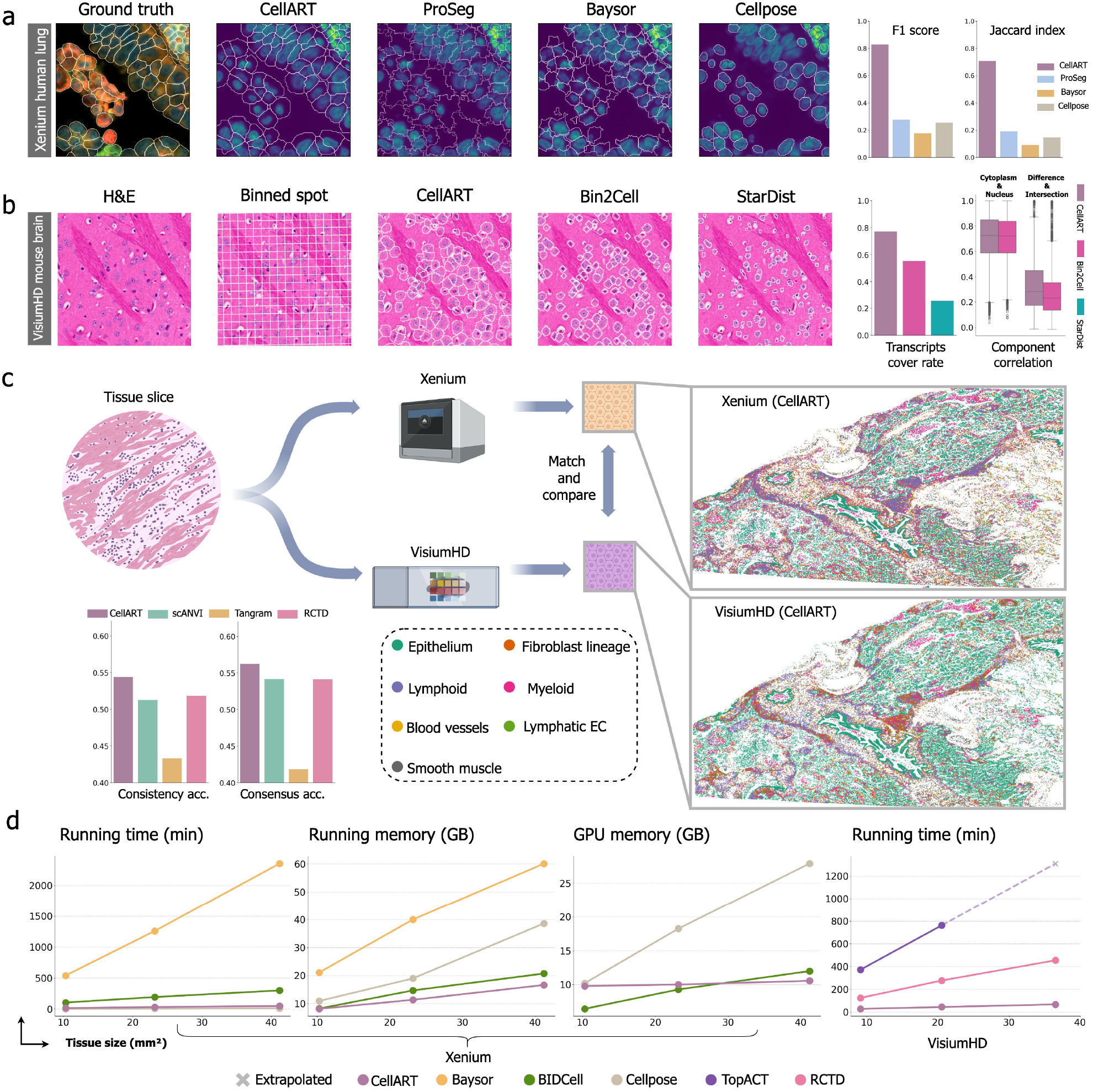
Performance evaluation of various methods on the Xenium and VisiumHD datasets. **a.** Comparison of cell segmentation results from CellART, ProSeg, Baysor, and Cellpose against the ground truth on the Xenium 2.0 dataset of human lung tissue. The ground truth is derived from the mixed-fluorescence image provided by 10x Genomics. The F1-score and Jaccard index are used to quantify segmentation accuracy. **b**. Comparison of cell segmentation results from CellART, Bin2cell, and StarDist on the VisiumHD Mouse Brain dataset. The analysis includes 16 *µ*m binned spots recommended by 10x Genomics, which do not align well with the cells in the histology image. The transcript coverage rate is computed to assess performance. Additionally, we compare CellART with Bin2cell using Pearson correlation, calculated between: 1. the cytoplasmic gene expression profile and the nuclear gene expression profile for each cell; and 2. the expression in different areas versus the intersection area of CellART and Bin2cell segmentation results for each cell. **c**. Validation of cell type label recovery using a paired human lung tissue dataset of a same tissue profiled with Xenium and post-Xenium VisiumHD, demonstrating CellART’s superior annotation accuracy and robustness across spatial transcriptomics platforms. **d**. Benchmarking computational efficiency across different methods. The Xenium and VisiumHD datasets of varying tissue sizes (x-axis) are created by subsampling the original datasets.

We start by evaluating the accuracy of cell boundary recovery on a Xenium Human Lung Cancer dataset (Fig. 2**a**, Supplementary Fig. S8), which contains 162,254 cells and 377 genes defined by 10x Genomics [28], with a median transcript count of 46 per cell. The dataset also provides cell segmentation masks derived from multiple morphological staining images which we regard as ground truth for evaluation. CellART successfully detected 156,335 cells, achieving a median transcript count of 52 per cell, closely matching the ground truth. The small number of undetected cells primarily resulted from incomplete nuclear staining, which hindered their identification (Supplementary Fig. S4); a dedicated preprocessing pipeline for nuclei quality control and a representation-based missing cell detection method are provided to diagnose and mitigate such cases (Supplementary Figs. S40 and S41). In addition, the cell boundaries predicted by CellART exhibited precise alignment with the ground truth, whereas ProSeg [21] and Baysor [22] produced substantially more segmentation artifacts (Fig. 2**a**). CellART also effectively delineated cytoplasmic regions that image-only methods, such as Cellpose [19], were unable to capture. Quantitative metrics further validated CellART’s superior performance, yielding markedly higher F1-scores (0.81) and Jaccard index values (0.70) than existing methods. The key to CellART’s success lies in its effective integration of transcriptomic and imaging data, which is superior to alternative methods that rely on a single modality.

Next, we assessed CellART’s performance on sequencing-based ST data using the VisiumHD Mouse Brain dataset [29] (Fig. 2**b**, Supplementary Fig. S9). This dataset lacks ground truth segmentation masks since the commonly used 16 *µ*m binned spots do not correspond to true cell boundaries. CellART processed raw 2 *µ*m subcellular spots, segmenting 61,851 cells. It provided single-cell resolution with accurate cell boundaries, outperforming Bin2Cell [18] and StarDist [20] by achieving higher transcript coverage rates and better capturing diverse cell morphologies. Specifically, CellART achieved a transcript coverage rate of 77%, compared to only 25% for StarDist. Two correlation-based metrics were used to evaluate segmentation quality (Supplementary Fig. S5, Methods). The first metric measured the consistency of gene expression between nuclear and cytoplasmic regions. The second metric evaluated the correlations between intersecting and non-overlapping regions of the CellART and Bin2Cell segmentation masks. Intersecting regions represent areas confidently assigned to the same cell by both methods and are expected to exhibit expression profiles that align with the remaining non-overlapping parts of that cell. CellART consistently achieved higher correlations in both metrics compared to Bin2Cell, while StarDist was excluded from this analysis as it does not include cytoplasmic regions in its segmentation results. Furthermore, CellART maintained larger cell sizes, which aligns with biological expectations and demonstrates its ability to accurately assign transcripts to cells. This superior performance is attributed to CellART’s design, which predicts cell boundaries using high-resolution representations that capture transcriptomic similarities between subcellular spots. Since the subcellular spots in the cytoplasm and nucleus naturally share similar transcriptomic profiles, this approach enables highly accurate boundary delineation. In contrast, Bin2Cell employs a simpler strategy of nuclei dilation to define cell boundaries, which often results in suboptimal segmentation outcomes. In summary, CellART not only captures more transcripts than existing methods, but also accurately assigns them to cells.

To validate the cell type annotation, we utilized a dataset [30] where a slice of human lung tissue was analyzed using both Xenium and post-Xenium VisiumHD profiling (Fig. 2**c**). We aligned the results obtained from the two technologies to establish a robust comparative framework and compared the cell type labels assigned to the same cells across platforms. Since only CellART can jointly perform segmentation and annotation, we used the Xenium segmentation provided by 10x Genomics and the StarDist-segmented VisiumHD cells for other annotation methods. After filtering out unmatched cells, 142,782 cells were retained for comparison. Annotation consistency between the two platforms revealed that CellART outperformed existing approaches such as scANVI, Tangram and RCTD, effectively addressing challenges such as platform effects and low sensitivity. CellART demonstrated consistent cell type distributions across both platforms (Fig. 2**c**), highlighting its robustness in handling both imaging-based and sequencing-based ST data. To ensure a fair and robust benchmark, we used Xenium-derived consensus labels as the ground truth for the comparison. These consensus labels were generated through a majority-vote strategy across all tested methods, requiring agreement from at least three of the four methods, thus providing reliable labels for comparison. The rationale for using these labels is twofold: (1) Xenium is performed prior to post-Xenium VisiumHD and thus does not influence the data quality, offering higher sensitivity; and (2) the majority-vote consensus ensures robust and biologically meaningful annotations, addressing the lack of real ground truth in spatial transcriptomics data. We note that these consensus labels represent surrogate ground truth rather than experimentally verified cell type identities, which could be a limitation of benchmarking cell type annotation in spatial transcriptomics where true ground truth labels are rarely available. Using these consensus labels, CellART demonstrated superior annotation accuracy on VisiumHD, further validating the reliability and precision of the recovered cell type labels. The high degree of agreement between CellART’s annotations on Xenium and VisiumHD highlights its robustness and the ability to accurately annotate cells from both imaging-based and sequencing-based platforms.

Scalability and computational efficiency are essential for handling large-scale, high-resolution ST datasets, which often contain tens of millions of transcripts and image data exceeding hundreds of gigabytes in size. We benchmarked CellART on Xenium and VisiumHD datasets of varying tissue sizes (Supplementary Fig. S7), demonstrated its ability to handle large-scale data with exceptional efficiency (Fig. 2**d**). The efficiency of CellART arises from its model architecture, which incorporates a patchification strategy to minimize memory requirements (Methods). Additionally, the lightweight and highly optimized neural network design further reduces computational costs. These innovations enable CellART to maintain high efficiency in both time and memory usage, completing the analysis of full-size datasets within three hours for both platforms while requiring less than 12 GB of GPU memory for Xenium datasets. In contrast, Baysor exhibited significant increases in runtime with larger tissue sizes, while Cellpose encountered memory limitations that constrained its scalability. In addition to annotation and segmentation methods, we also observed that TopACT [16], a method specifically designed for direct cell type annotation on subcellular spots, lacks scalability to whole-tissue datasets. The ability of CellART to process large-scale data with high efficiency establishes its practical applicability for large-scale studies.

In summary, our benchmarking study demonstrates CellART’s outstanding performance in extracting single-cell information from high-resolution ST datasets. It excels in recovering precise cell boundaries and accurate cell type labels, while maintaining high computational efficiency, making it a practical tool for high-resolution ST data.

### CellART demonstrates robustness and versatility in cross-platform validation using mouse brain datasets

To evaluate the generalizability of CellART across diverse platforms, we applied it to mouse brain datasets generated using four distinct high-resolution ST platforms: VisiumHD [29], Xenium [31], MERFISH [32], and Stereo-seq [33]. These datasets differ in spatial resolution, gene coverage, and experimental protocols, offering a rigorous test of CellART’s adaptability and robustness.

We used a single scRNA-seq dataset from the Allen Mouse Brain Atlas[34], which includes 40 cell types, as the reference for all ST datasets to ensure consistency in cell type labeling (Supplementary Fig. S10). The results from CellART (Fig. 3**a, b**) demonstrate a high degree of concordance in cell type annotations, despite variations in technical specifications across platforms. CellART accurately identified major cell types, including neuronal and glial populations, and reconstructed their spatial distributions with high fidelity. Cell-level statistics revealed technical variations. For instance, we utilized the whole gene panel of Xenium, comprising 248 genes, whereas for VisiumHD, we first identified 2,000 highly variable genes from the scRNA-seq reference and used these genes as input for CellART to analyze the VisiumHD data. Consequently, CellART successfully identified 161,762 cells in the Xenium mouse brain dataset and 61,851 cells in the VisiumHD mouse brain dataset. Although Xenium has fewer genes in its panel, the median number of transcripts per cell in Xenium was 172, significantly higher than the median of 65 transcripts per cell observed in VisiumHD, reflecting the high sparsity of transcripts in VisiumHD. Despite these differences, the reconstructed cell sizes were comparable, and the relative abundances of the ten most prevalent cell types, such as astrocytes and oligodendrocytes, were highly consistent across the two platforms (Fig. 3**c**). CellART’s annotation for the MERFISH and Stereo-seq datasets further revealed intricate spatial structures that aligned closely with known biological patterns, demonstrating consistency across all platforms (Supplementary Fig. S11, S12). Moreover, CellART consistently and accurately reconstructed the cortical lamination, including layers L2/3, L4, L5, and L6 (Fig. 3**d**), across all four platforms, closely matching known anatomical landmarks and further demonstrating its robustness and adaptability across different technologies.

**Figure 3.**
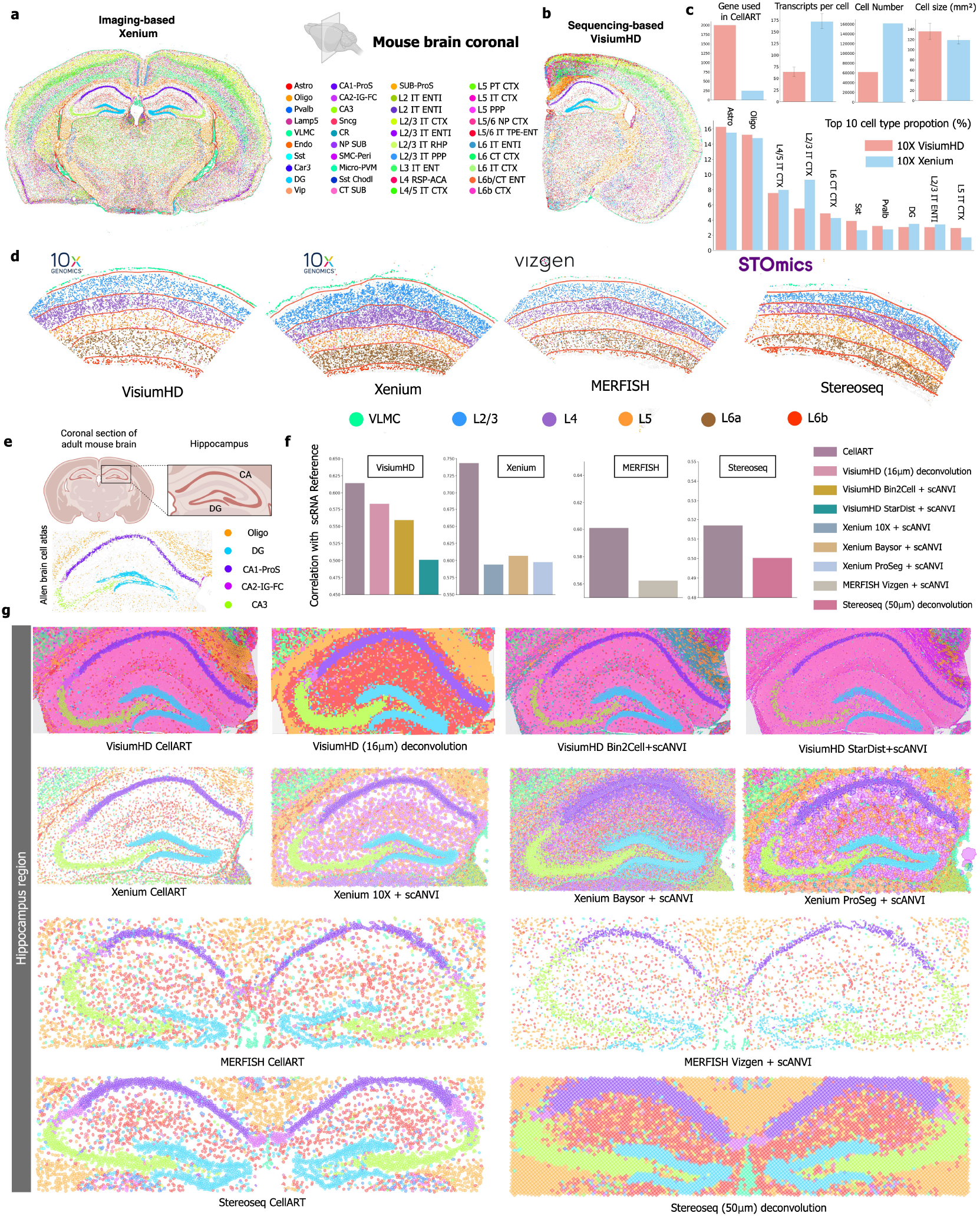
Application of CellART to mouse brain datasets from different platforms. **a.** Cell annotation results obtained from CellART on the Xenium mouse brain dataset. **b**. Cell annotation results obtained from CellART on the VisiumHD mouse brain dataset. **c**. Barplots of different summary statistics of cells extracted by CellART on Xenium and VisiumHD mouse brain datasets. On the top is the number of genes used, average UMI, cell number and cell size. Below are the proportions of the ten most abundant cell types, showing highly consistent results between the two platforms. **d**. Reconstructed layer structure of the mouse brain on VisiumHD, Xenium, MERFISH, and Stereo-seq datasets using CellART. Red curves indicate the approximate layer borders. **e**. Annotation from Allen Mouse Brain reference of the hippocampus region. **f**. Pearson correlation between the extracted cells and the single-cell reference data for each method. **g**. Visualization of cell segmentation and annotation results obtained from CellART and other methods on VisiumHD, Xenium, MERFISH and Stereo-seq mouse brain datasets in the hippocampus region.

In brain tissue, the hippocampus plays a crucial role in learning and memory [35]. Its anatomical position and reference spatial annotations [36] from the Allen Mouse Brain Atlas are shown in Fig. 3**e**, highlighting major hippocampal subregions, including the Cornu Ammonis areas (CA1, CA2, CA3), the dentate gyrus (DG), and surrounding oligodendrocytes (Oligo). CellART demonstrates high precision in segmenting and annotating these subregions, achieving the best alignment of annotations with reference spatial data. In contrast, baseline methods exhibited significant limitations (Fig. 3**g**). For the VisiumHD dataset, deconvolution-based approaches were unable to achieve single-cell resolution, resulting in an inability to accurately delineate hippocampal structures. Since Bin2Cell and StarDist perform segmentation and scANVI [37] handles annotation, we combined them as Bin2Cell+scANVI and StarDist+scANVI as additional baselines for the VisiumHD dataset. The results shows that these combinations introduced errors, including oligodendrocyte misclassification and mixing of CA cell types. For the Xenium dataset, the 10x Genomics segmentation overestimated cell sizes, whereas Baysor and ProSeg generated artifact-prone segmentations due to their lack of integration with staining images, resulting in unrealistically high cell densities. Additionally, scANVI, when applied to cells segmented by baseline methods, failed to correctly identify the CA2 region in the Xenium dataset. For the MERFISH dataset, CellART provided more complete and biologically coherent segmentation compared to the official Vizgen segmentation, which relied solely on DAPI staining. CellART achieved higher transcript coverage and better alignment with the spatial annotation reference in Fig. 3**e**. When scANVI was applied to Vizgen-segmented cells, it again failed to recover CA2 structures. In the Stereo-seq dataset, where conventional methods such as RCTD [14] operate at multicellular resolution using 50 *µ*m bins, CellART achieved true single-cell resolution. These results underscore CellART’s superior ability to extract single-cell level information from both imaging-based and sequencing-based spatial transcriptomics platforms. Furthermore, CellART achieved the highest Pearson correlation (Methods) with single-cell reference data across all platforms (Fig. 3**f**), confirming its accuracy and reliability.

Additional validation was provided by marker gene analysis in the hippocampus (Supplementary Fig. S13). Violin plots of canonical hippocampal markers, including *Prox1* (DG), *Necab2* (CA2), *Slit2* (CA3), and *Wfs1* (CA1), confirmed that CellART accurately identified neuronal subtypes and recovered their spatial locations consistent with previous studies. Notably, CellART also identified *Cpne8* as a distinct gene enriched in CA1, further demonstrating its ability to uncover biological signals. The spatial localization of these markers further verified that CellART faithfully recovered cellular architecture aligned with reference hippocampal anatomy.

Having established CellART’s methodological performance—including segmentation accuracy, annotation precision, computational efficiency, and cross-platform generalizability—we next turn to two cancer case studies to demonstrate the biological insights that CellART’s unified framework and enhanced single-cell resolution can enable. In these applications, methodological comparisons with baseline methods are presented alongside biological analyses, as the advantages of CellART are most apparent in the challenging tissue contexts where they matter most.

### CellART accurately extracts cellular information and identifies transient cancer cells in Xenium breast cancer datasets

Breast cancer is the most common malignancy in women and exhibits pronounced intratumoral heterogeneity, where transient and intermediate cellular states shape invasion, therapy response, and relapse. To interrogate these states at single-cell resolution and in their native spatial context, we applied CellART to a Xenium breast cancer dataset [7] profiled with a targeted panel of 313 genes. Leveraging its unified segmentation-annotation framework, CellART delineated precise cell boundaries, resolved tumor and stromal compartments, and revealed spatial gene programs and subtype dynamics.

The recovery of cell type labels (Fig. 4**a**) demonstrates CellART’s capacity to resolve distinct cell types at high spatial fidelity. To assess the replicability of CellART, we compared the results between two adjacent slices (Xenium breast cancer replicate 1 and Xenium breast cancer replicate 2). The annotation results from both slices were highly consistent, demonstrating that CellART is robust and replicable. This reproducibility is further supported by the UMAP plot (Supplementary Fig. S14), which shows a well-mixed population of cells between the two replicate datasets. CellART also successfully identified and accurately assigned markers to cell types, such as *EPCAM, FOXA1*, and *ERBB2* for breast cancer cells, and LUM for stromal cells (Supplementary Fig. S17).

**Figure 4.**
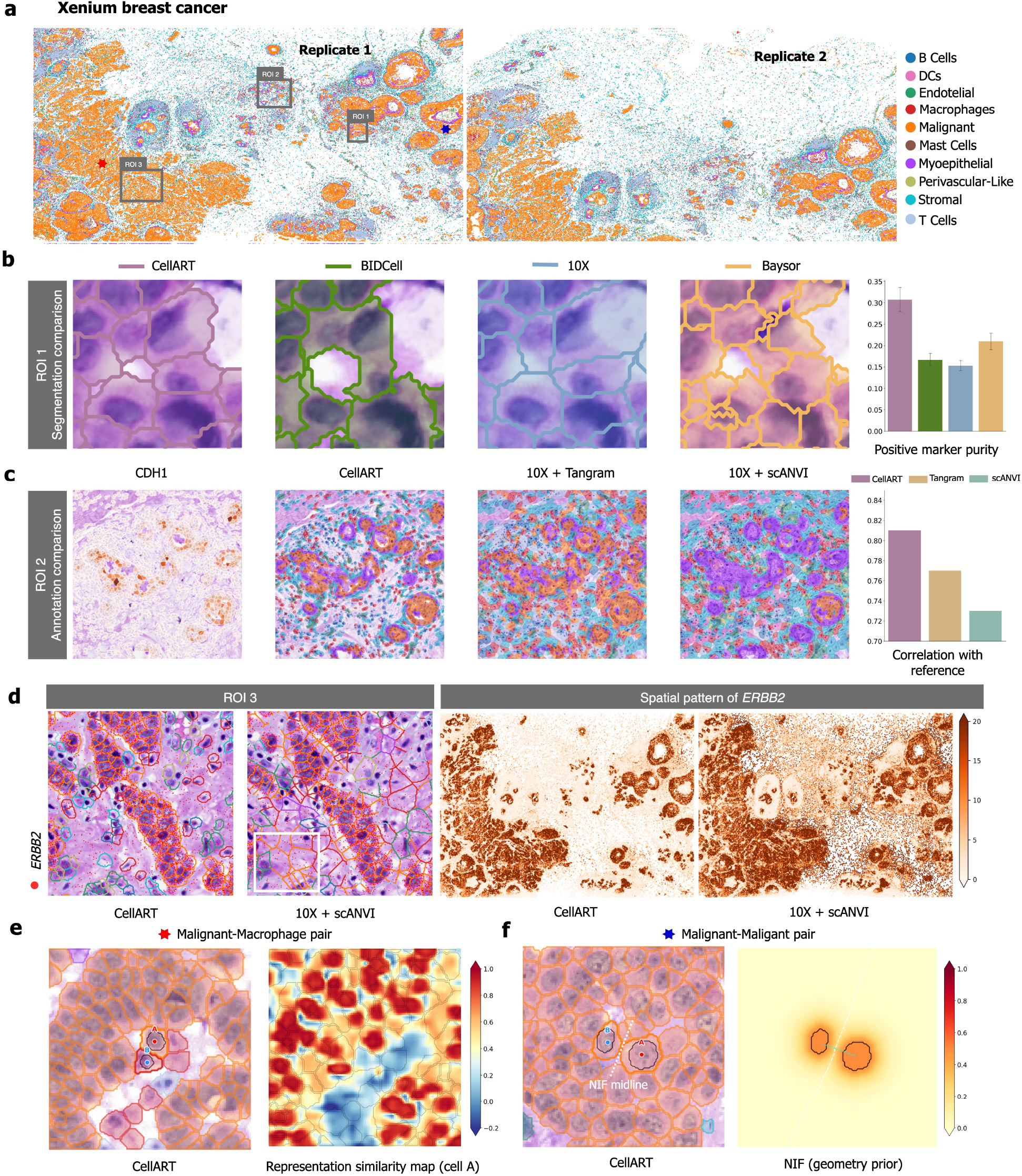
Application of CellART to the Xenium breast cancer dataset. **a.** Visualization of cell type annotations from the Xenium breast cancer dataset across replicates 1 and 2. CellART accurately resolves diverse cell types, showing a high consistency between replicates. Three regions of interest (ROIs) are highlighted for detailed comparison to validate performance across distinct regions and cell types. **b**.Segmentation comparison within ROI-1 between CellART, BIDCell, 10x, and Baysor. CellART achieves precise cell boundary delineation, avoiding nucleus merging (BIDCell) and fragmented boundaries (Baysor). Quantitative evaluation using positive marker purity confirms CellART’s superior segmentation accuracy, achieving the highest purity. **c**. Annotation comparison in ROI-2. CellART successfully identifies tumor clusters that align closely with *CDH1* expression. In contrast, Tangram introduces false-positive tumor annotations, and scANVI fails to distinguish between tumor and myoepithelial cells. Pearson correlation analysis with scRNA-seq references validates CellART’s higher annotation accuracy. **d**.Spatial distribution of *ERBB2* expression in ROI-3 and across the dataset. CellART delineates sharp and biologically accurate *ERBB2* expression patterns while minimizing transcript diffusion. In contrast, 10x introduces noise and misaligned transcripts. **e**. Adaptive boundary behavior between transcriptionally distinct cell types (malignant–macrophage pair). Left: CellART segmentation with the focal cell pair highlighted. Right: representation similarity map to cell A’s nucleus, showing a clear gradient that distinguishes the two cell types and confirms expression-driven boundary placement. **f**. Adaptive boundary behavior between transcriptionally identical cell types (malignant–malignant pair). Left: CellART segmentation with the NIF midline (white dashed). Right: NIF geometric prior, showing that under transcriptional homogeneity the boundary aligns with the NIF equal-influence contour, producing a Voronoi-like tessellation.

For cell segmentation (Fig. 4**b**, Supplementary Fig. S15**a**), CellART demonstrates a significant advantage over baseline methods by reliably capturing individual cell boundaries. BIDCell [38] often merged multiple nuclei into a single cell, while CellART accurately separated them. Additionally, the segmentation produced by CellART closely aligned with regions of deep staining that indicate cellular presence, while avoiding the inclusion of blank areas, a recurring issue observed in the 10x segmentation approach. In contrast, Baysor generated fragmented boundaries that frequently intersected nuclei, which could be biologically misleading. To assess the accuracy of cell segmentation in the absence of ground truth, we employed the positive marker purity metric (Methods). This metric leverages marker genes derived from scRNA-seq reference data, where positive markers refer to the top 10% of highly expressed genes present in at least 50% of cells for each cell type. Alternative segmentation methods were paired with scANVI for annotation, whereas CellART simultaneously predicted both cell boundaries and cell type labels. For each cell, the sensitivity was calculated as the proportion of positive markers expressed within the cell, with box plots used to visualize the distribution of sensitivity across all cells. High positive marker purity indicates that a segmentation method accurately reflects the gene expression profiles of the respective cell types. CellART achieved the highest positive marker purity among all tested methods, demonstrating its ability to produce reliable segmentation and biologically consistent cell annotations. Furthermore, CellART excels at preserving cell morphology. By calculating the real nuclei eccentricity from staining images and comparing it to segmented cell eccentricity, CellART achieved a correlation as high as 0.94, BIDCell showed a comparable correlation as it uses prior knowledge of cell type elongation. In contrast, Baysor and 10x yielded correlations below 0.5, reflecting their inability to preserve diverse cell morphologies (Supplementary Fig. S24). Descriptive statistics further highlight CellART’s ability to balance cell size and transcriptional density, maintaining moderate cell sizes while achieving high transcript density, ensuring biologically meaningful segmentation and annotation (Supplementary Fig. S16).

Cell type annotation with CellART also enables robust tumor cell identification (Fig. 4**c**, Supplementary Fig. S15**b**). Tumor clusters identified by CellART aligned closely with the distribution of the cancer marker *CDH1*. In comparison, Tangram [39] introduced false-positive tumor cells in regions outside true tumor clusters, while scANVI was unable to distinguish between cancer and myoepithelial cells. To quantitatively assess annotation accuracy, we calculated the Pearson correlation coefficient between the average gene expression profile of annotated cells for each cell type and the corresponding cell type in the scRNA-seq reference. These correlation coefficients were then averaged across cell types. CellART achieved an average correlation exceeding 0.8, higher than other methods, showing its ability to recover biologically accurate cell type annotations. Additionally, CellART showed consistent proportions of annotated cell types when compared with scRNA-seq references derived from the same tissue (Supplementary Fig. S23).

To highlight CellART’s ability in precisely delineating cell boundaries, we performed a detailed analysis of the spatial expression distribution of *ERBB2*, a key tumor marker encoding a receptor tyrosine kinase in the *EGFR* family (Fig. 4**d**). Unlike the 10x standard pipeline, which often retains noisy transcripts, CellART produces clean and accurate spatial expression patterns of *ERBB2* that closely align with tumor cell distributions. Furthermore, cell type distributions inferred by CellART align well with their corresponding marker gene distributions (Supplementary Fig. S17). To illustrate how CellART adaptively determines cell boundaries, we examined two contrasting scenarios (Fig.4**e,f**). In cases where neighboring cells are transcriptionally distinct, such as a malignant-macrophage pair, boundary placement is guided by learned representation similarity, resulting in a sharp gradient at the cell type interface (Fig.4**e**). In contrast, when neighboring cells share similar transcriptional profiles, such as two adjacent malignant cells, their boundaries are primarily influenced by the NIF geometric prior, leading to a Voronoi-like tessellation that aligns with the NIF midline (Fig. 4**f**). Extended diagnostics, including boundary pixel gene expression analysis, segmentation uncertainty, and annotation uncertainty visualization, are provided in Supplementary Figs. S44–S46. Niche analysis and ligand-receptor interactions between myoepithelial and tumor cells are presented in Supplementary Figs. S19, S20, and S22.

CellART further resolves tumor subtypes with single-cell and subcellular precision (Supplementary Fig. S39). Existing segmentation methods lack the ability to naturally pair nuclei with cytoplasm, which limits their precision. For example, methods like BIDCell often include multiple nuclei within a single boundary, while others such as Baysor and ProSeg produce fragmented or biologically implausible boundaries that cross nuclei. In contrast, CellART’s framework incorporates image data to precisely define the nuclei and uses them as anchors to extend to accurate cell boundaries based on transcriptomic similarity (Methods). This principle design yields a one-to-one nucleus-cell correspondence and a faithful partition of nuclear and cytoplasmic compartments, enabling downstream subcellular analyses that are otherwise infeasible. Leveraging these accurate boundaries, CellART robustly recapitulates tumor subtypes, including proliferative invasive tumor, invasive tumor, DCIS 1, and DCIS 2, consistent with pathologist annotations from the original study [7] (Supplementary Fig. S21). CellART’s precise nucleus-cytoplasm pairing also enables exploratory RNA velocity analysis by treating cytoplasmic transcripts as mature and nuclear transcripts as nascent RNA (Supplementary Fig. S39). These velocity results should be interpreted as hypothesis-generating, as nuclear localization does not always correspond cleanly to transcriptional state.

### CellART enhances resolution and reduces transcript misallocation in VisiumHD colorectal cancer

Colorectal cancer is a leading cause of cancer-related mortality and exhibits pronounced spatial heterogeneity, where boundary doublets and transcript mixing obscure true cellular states and interactions. To evaluate whether CellART can enhance VisiumHD analyses by advancing from binned-spot deconvolution to true single-cell resolution and minimizing transcript misallocation, we analyzed a VisiumHD colorectal cancer dataset [3] at 2 *µ*m raw-spot resolution and leveraged a paired Xenium dataset from the same patient for cross-platform validation. Conventional analyses of VisiumHD data rely on 8 *µ*m or 16 *µ*m binned spots, typically processed using deconvolution methods such as RCTD to infer cell types. However, this strategy often causes transcript mixing, particularly in doublet spots at domain boundaries, thereby obscuring cellular identities (Supplementary Fig. S25). CellART addresses these limitations by directly extracting single-cell information from raw subcellular spots, providing a more detailed and biologically accurate representation of cellular spatial organization.

CellART successfully resolved individual cell types at single-cell resolution, with annotations demonstrating high concordance with the Xenium colorectal cancer dataset derived from the same patient, underscoring its robustness and cross-platform consistency (Fig. 5**a**). Using the scRNA-seq reference from the same patient, CellART identified 256,238 cells across 28 cell types in the VisiumHD colorectal cancer dataset. A comparison with the 8 *µ*m deconvolution result from RCTD from the original study further highlights CellART’s advantages. In ROI-1, CellART transitioned from the coarse resolution of 8 *µ*m binned spots to precise single-cell annotations, revealing spatially distinct cell types blurred in conventional binned analyses (Fig. 5**b**). This enhanced resolution allowed for the accurate identification of individual cells, even in densely packed tumor and immune cell regions (Supplementary Fig. S30). Marker gene expression analysis further validated CellART’s segmentation and annotation, with tumor markers such as *CEACAM6* and *AXIN2* accurately localized within tumor cells, demonstrating the biological accuracy of the reconstructed spatial organization (Supplementary Fig. S26).

**Figure 5.**
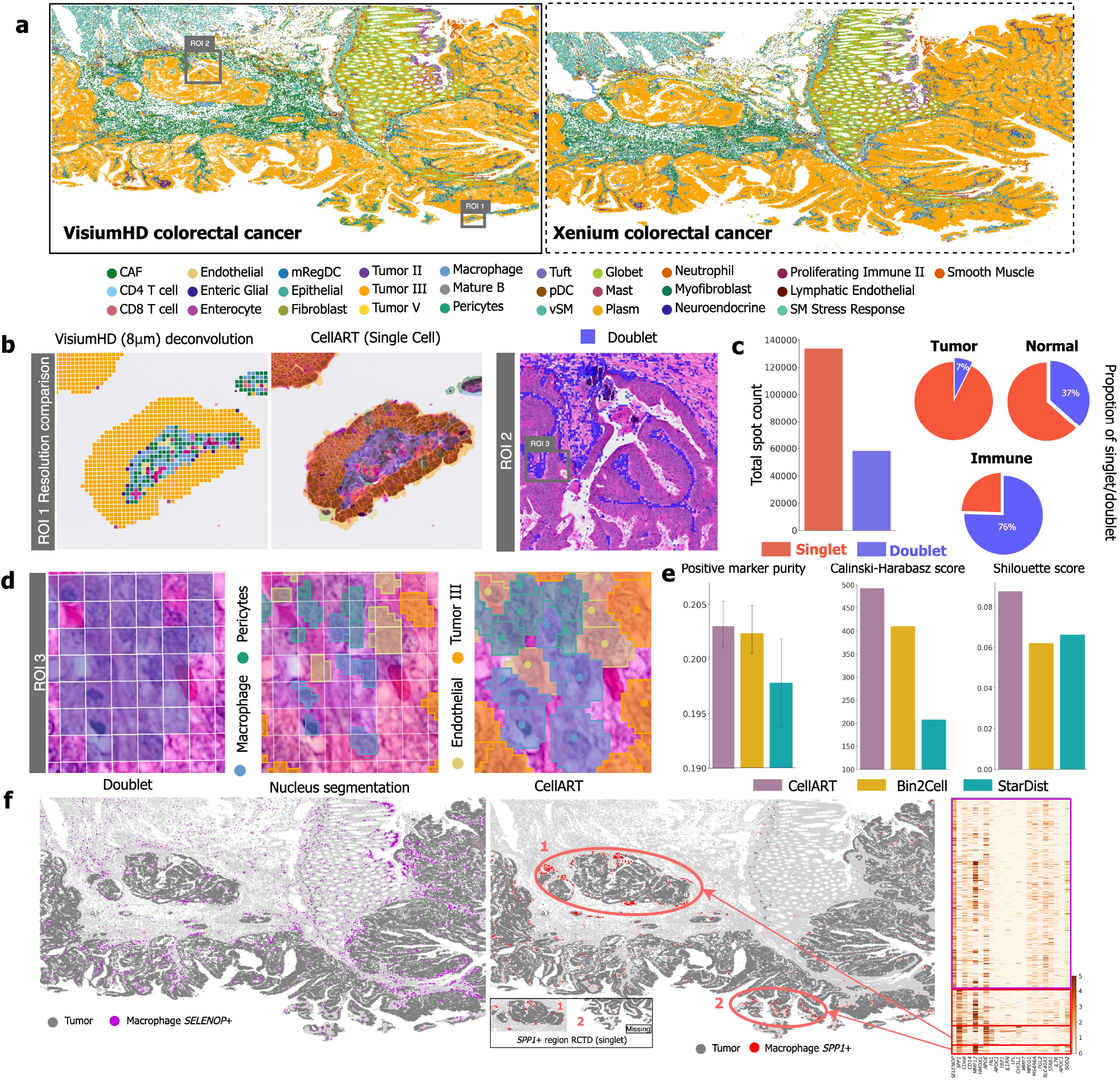
Application of CellART to the VisiumHD colorectal cancer dataset. **a.** Cell type annotations by CellART on the VisiumHD colorectal cancer dataset, demonstrating its ability to resolve individual cell types at single-cell resolution. The annotations are compared with those from the Xenium colorectal cancer dataset of the same patient, showing high consistency across platforms. Two regions of interest (ROIs) are highlighted for detailed comparisons. **b**.ROI-1: Resolution comparison between RCTD deconvolution of 8 *µ*m binned spots and CellART’s single-cell segmentation. CellART significantly enhances resolution, transitioning from coarse binned spots to precise single-cell annotations. ROI-2: Visualization of a doublet region, where RCTD identifies doublet spots (blue), predominantly located at the boundaries of tumor cell clusters. **c**. Quantitative assessment of RCTD’s deconvolution results. Left: A bar plot showing the proportion of singlet and doublet spots, revealing that approximately one-third of the spots are classified as doublets. Right: Pie charts illustrating the distribution of spot classes for tumor, immune, and other cells. Tumor cells, densely packed, exhibit the lowest proportion of doublets, while immune cells, located at tumor boundaries and including diverse types like CAFs and macrophages, show over 75% doublets. **d**. Detailed analysis of a doublet region in ROI-2. Left: Doublet spots (blue) identified by RCTD. Middle: Nuclei segmentation results from StarDist, with grid lines representing spot borders, showing nuclei crossing spot boundaries. Right: CellART’s segmentation results, recovering accurate cell boundaries and assigning cell type labels. **e**. Quantitative comparison of CellART, Bin2Cell, and StarDist based on marker gene purity, Calinski-Harabasz index, and Silhouette score. CellART consistently achieves higher marker purity and superior clustering metrics, indicating better segmentation quality. RCTD is excluded from this comparison as it operates at spot-level deconvolution rather than single-cell segmentation, making direct metric comparison inequitable. **f**. Spatial distribution of *SELEONOP* + and *SPP1* + macrophages identified in the original study. Heatmap (right) reveals expression profiles of these macrophage subtypes. Two *SPP1* + macrophage regions identified by CellART are highlighted (red circles). Region-1 corresponds to singlet regions recognized by RCTD, while region-2 is uniquely detected by CellART’s improved resolution, revealing previously obscured macrophage subpopulations.

A key challenge in 8 *µ*m binned spot-level analyses is the prevalence of doublet spots, particularly along tumor boundaries, where cellular mixing is high due to the complex microenvironment enriched with diverse cell types such as immune cells, cancer-associated fibroblasts (CAFs), and tumor cells. ROI-2 provides a closer examination of this phenomenon, showing that doublets predominantly occur at tumor boundaries (Fig. 5**b**). Approximately one-third of all 8 *µ*m binned spots in the dataset were identified as doublets by RCTD, with immune-cell-majority spots exhibiting the highest doublet rate (76%) due to their frequent proximity to tumor clusters and the intricate interactions at the tumor’s periphery (Fig. 5**c**). These doublets obscure cell type-specific transcript signals, leading to annotation inaccuracies and limiting the resolution of downstream analyses. CellART effectively resolved this issue by recovering individual cells within such regions, accurately segmenting cell boundaries and assigning precise cell type labels. In ROI-3, a zoomed-in region of ROI-2, most spots were classified as doublets (blue) (Fig. 5**d**). Nuclei segmentation results obtained from StarDist, using H&E staining, revealed that spot boundaries often crossed multiple nuclei, each corresponding to different cell types. In contrast, CellART reconstructed entire cell boundaries, resolving transcript misallocation due to segmentation inaccuracies and providing clear assignments of cell identities. Quantitative benchmarking against alternative single-cell methods, including Bin2Cell and StarDist, confirmed CellART’s superior performance, achieving the highest positive marker purity, Calinski-Harabasz score, and Silhouette score (Fig. 5**e**). RCTD was excluded from this quantitative comparison because it operates at spot-level deconvolution rather than single-cell segmentation, making direct metric comparison inequitable; instead, its relationship to CellART is better understood as complementary, with RCTD’s doublet calls faithfully reflecting the ambiguity inherent to the 8 *µ*m binned-spot geometry.

To further elucidate the biological insights enabled by CellART, we focused on analyzing macrophage populations, which play a critical role in tumor-immune interactions. At the 8 *µ*m spot level, RCTD correctly identifies macrophage-majority spots as doublets when they capture transcripts from multiple cell types at tumor boundaries (Supplementary Fig. S31), reflecting genuine spatial ambiguity inherent to the binned-spot geometry. CellART’s single-cell resolution complements this by disentangling the mixed signals into individual cells, enabling the identification of immune-specific markers such as *CD4* and *CD53* within macrophage populations. Differential expression analysis comparing macrophage populations identified by RCTD (doublet spots) and CellART (segmented single cells) is presented in the Supplementary Fig. S28, demonstrating that CellART’s single-cell resolution recovers immune-specific transcriptional programs that are diluted in spot-level analyses.

Beyond resolving conventional cell types, CellART revealed cellular subpopulations with distinct functional roles. Notably, the *SELEONOP* + and *SPP1* + macrophage subtypes, originally identified in the reference study, were also detected by CellART, each exhibiting distinct transcriptional signatures (Fig. 5**f**). Furthermore, CellART’s single-cell resolution enabled the discovery of two spatially and molecularly distinct *SPP1* + macrophage regions. Region-1, recognized by both RCTD and CellART, was located near tumor boundaries and characterized by enrichment of genes such as *APOE* and *FN1* (Supplementary Fig. S27), implicating these macrophages in local tumor-immune crosstalk. In contrast, region-2, uniquely detected by CellART, revealed macrophage subpopulations that were previously obscured due to the limitations of spot-based analyses. Macrophages in region-2 were enriched for genes like *MMP12* and *HMOX1*, which are associated with remodeling the extracellular matrix (ECM) and promoting cancer invasion [40]. This molecular divergence was mirrored by the behavior of adjacent cancer cells: those in region-2 exhibited significantly reduced *CDH1* (E-cadherin) expression compared to region-1 (*t*-test, *p <* 0.001; Supplementary Fig. S27), a hallmark of enhanced migratory and invasive potential [41], suggesting that the tumor cells in region-2 are more likely to migrate and invade surrounding tissues. By resolving these subtle macrophage subpopulations and linking their molecular states to cancer cell behaviors, CellART demonstrates its ability to overcome the limitations of conventional spot-based methods and provide critical mechanistic insights into the spatial orchestration of the tumor microenvironment.

Cell niche analysis using CellART annotations is presented in Supplementary Fig. S32, where 11 biologically meaningful niches were identified with high consistency between the Xenium and VisiumHD datasets. Among these, macrophage-enriched niches adjacent to tumor regions motivated the following investigation into tumor-associated macrophages.

### CellART reveals consistent tumor-associated macrophages in VisiumHD and Xenium colorectal cancer

Building on CellART’s ability to achieve single-cell resolution and mitigate transcript misallocation in the VisiumHD colorectal cancer dataset, we leveraged its precision to identify TAMs and investigate their spatial and molecular characteristics. As a functionally distinct subset of macrophages, TAMs were defined as those residing within 50 *µ*m of tumor cells, indicating their spatial proximity and biological relevance within the tumor microenvironment (TME). The investigation began with characterizing the spatial distribution and distinct expression profiles of TAMs in VisiumHD, while extending the analysis to the Xenium colorectal cancer dataset of the same patient to confirm consistency. Then, TAM-tumor communication was explored to uncover key biological insights.

The spatial distribution of TAMs, macrophages, and tumor cells was visualized across the tissue, revealing a pronounced enrichment of TAMs along tumor boundaries (Fig. 6**a**). CellART provided a refined view of TAM-tumor associations, resolving individual cell identities and their spatial interactions. To highlight the advantages of CellART’s single-cell resolution, we focused on two TAM-enriched regions (ROI-1 and ROI-2). Within ROI-1, comparative analysis demonstrated that CellART accurately segmented TAMs and tumor cells, delineating their spatial boundaries with high precision (Fig. 6**b**). The corresponding region in the Xenium colorectal cancer dataset from the same patient showed high consistency with the VisiumHD, confirming the robustness of CellART across different platforms. A closer examination of ROI-2 revealed precise localization of TAM marker genes (Fig. 6**c**), such as *MMP12* and *SPP1*, within TAM-enriched regions identified by CellART. Notably, *SPP1* + macrophages identified in the previous analysis were confirmed to be part of the TAM population in VisiumHD. Gene expression profiling provided additional insights into the transcriptional specialization of TAMs. UMAP visualization revealed that TAMs formed a distinct cluster situated between tumor cells and normal macrophages, reflecting their hybrid transcriptional state induced by the tumor microenvironment (Fig. 6**d**). Heatmap analysis further confirmed that TAMs upregulated immune-related and tumor-promoting genes, especially *MMP12* compared to normal macrophages (Fig. 6**e**). These transcriptional signatures suggest that TAMs are functionally distinct, contributing to both immune modulation and tumor progression. Violin plots confirmed the elevated expression of *MYC* and *MMP12* in TAMs across both VisiumHD and Xenium datasets, showing the enrichment with high reliability. While *SPP1* expression, restricted to the VisiumHD dataset, highlighted the advantages of whole-transcriptome profiling in capturing TAM-specific markers (Fig. 6**f**).

**Figure 6.**
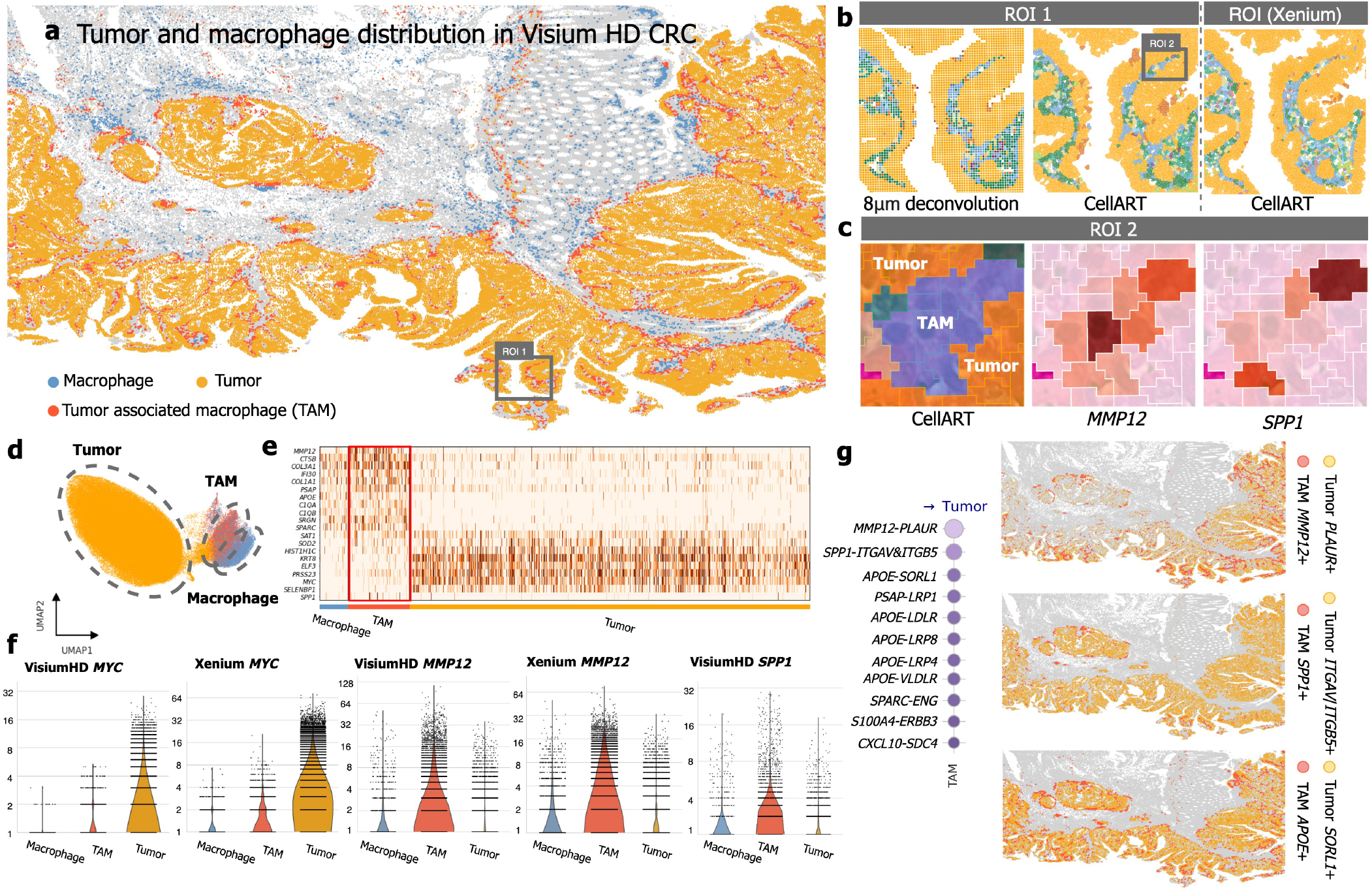
Identification of tumor-associated macrophages (TAMs) in the VisiumHD and Xenium colorectal cancer dataset. **a.** Spatial distribution of macrophages, tumors, and TAMs in the colorectal cancer microenvironment. TAMs are defined as macrophages with a minimum distance of less than 50 *µ*m from tumor cells. The visualization highlights the enrichment of TAMs along tumor boundaries, providing insights into their spatial association with the tumor microenvironment. **b**. ROI-1: Visualization of RCTD deconvolution of 8 *µ*m binned spots compared to single-cell resolution achieved by CellART, focusing on a TAM- and tumor-rich region. To validate the robustness of CellART’s annotations, we examined the same region in the Xenium dataset from the same patient. **c**. ROI-2: Detailed zoom-in of ROI-1 showing accurate segmentation and annotation of TAMs and tumors by CellART. Spatial expression patterns of TAM marker genes (*MMP12, SPP1*) match the localized TAMs extracted by CellART, validating the accuracy of its single-cell resolution. **d**. UMAP visualization of gene expression profiles for tumor cells, TAMs, and normal macrophages. TAMs form a distinct cluster situated between tumor cells and normal macrophages. **e**. Heatmap showing gene expression profiles of tumor cells, TAMs, and normal macrophages. TAMs exhibit enhanced expression of immune-related and tumor-associated genes, indicating their functional specialization within the tumor microenvironment. **f**. Violin plots of *MYC, MMP12*, and *SPP1* expression levels in macrophages, TAMs, and tumor cells. *MYC* and *MMP12* exhibit consistent expression patterns in both VisiumHD and paired Xenium datasets (recovered by CellART), while *SPP1* is only shown in the VisiumHD dataset as it is not included in the Xenium panel. **g**. Left: Magnitude of ligand-receptor interactions between TAMs and tumor cells, emphasizing the enriched communication between these two cell populations. Right: Spatial expression patterns of representative ligand-receptor pairs, including *MMP12* -*PLAUR, SPP1* -*ITGAV* &*ITGB5* and *APOE* -*SORL1*, showing strong co-localization of TAMs and tumor cells, consistent with their interaction.

These spatial and transcriptional insights underscore the potential interactions between TAMs and tumor cells, which are pivotal in shaping tumor-immune dynamics. Leveraging CellART’s cell-resolved outputs, we performed ligand-receptor analysis to interrogate the molecular dialogue between TAMs and tumor cells (Fig. 6**g**). Notably, key canonical ligand-receptor pairs such as *MMP12* -*PLAUR* and *SPP1* -*ITGAV* &*ITGB5* exhibited strong spatial co-localization. The interaction between *MMP12* and *PLAUR* (urokinase plasminogen activator receptor) enhances tumor cell invasion and metastasis within the CRC microenvironment [42]. The interaction between *SPP1* and the *ITGAV* &*ITGB5* heterodimer (integrin *α*v*β*5) on CRC cells activates downstream signaling pathways that enhance tumor progression, immune evasion, and resistance to therapy [43]. These findings emphasize the molecular pathways through which TAMs influence the tumor microenvironment, particularly in processes related to extracellular matrix remodeling and immune modulation. Interestingly, our analysis also identified the interaction of *APOE* -*SORL1*, a pathway previously associated with neurodegenerative diseases such as Alzheimer’s disease [44, 45]. Recent studies have implicated a neuron-specific role for *APOE* in immune response pathways [46, 47]. Meanwhile, *SORL1* has been implicated in breast cancer progression by regulating HER2 trafficking and distribution [48], as well as in promoting chemoresistance in ovarian cancer [49]. In colorectal cancer, our findings suggest that the *APOE* -*SORL1* interaction may similarly play a critical role, although the underlying mechanisms require further investigation. These results emphasize the broader applicability of CellART in uncovering novel molecular pathways, providing a foundation for future studies into TAM-associated signaling in cancer progression.

## Discussion

In this paper, we presented CellART, a unified framework that integrates staining images, spatial transcriptomics data, and scRNA-seq references through deep learning and probabilistic modeling to simultaneously perform cell segmentation and cell type annotation from high-resolution ST data. CellART demonstrated superior accuracy and computational efficiency across both imaging-based and sequencing-based platforms, and its applications to breast and colorectal cancer datasets highlighted the ability to uncover tumor cell state transitions, resolve macrophage subpopulations, and characterize tumor-immune interactions at single-cell resolution. For VisiumHD data in particular, CellART resolved transcript misallocation issues inherent to conventional binned-spot analyses, enabling biologically accurate spatial reconstruction.

CellART’s framework rests on four modeling assumptions. First, nuclei detected from staining images serve as spatial anchors for cell boundary recovery. In cases of incomplete nuclear staining, a representation-based method is employed to identify and address missing cells (see Supplementary Figs. S41). However, when nuclear staining is entirely absent, CellART cannot extract single-cell information, a limitation we acknowledge. Second, we assume a one-to-one correspondence between nuclei and cells, with edge cases like multinucleated cells managed through an optional multi-nuclei merge step (see Supplementary Fig. S40). Third, a Poisson likelihood model is utilized for transcript counts, and a systematic evaluation of four alternative distributions (NB, ZIP, ZINB, Tweedie) produced virtually identical results (see Supplementary Fig. S43). Lastly, scRNA-seq reference profiles are often needed for cell type annotation. CellART has demonstrated robust performance even in reference mismatch scenarios (see Supplementary Fig. S42), and a reference-free NMF mode offers an unsupervised alternative when suitable references are unavailable.

In its current form, CellART extracts single-cell information from two-dimensional, high-resolution spatial transcriptomics data across sequencing-based (VisiumHD, Stereo-seq) and imaging-based (Xenium, MERFISH) platforms. The unified framework provides a foundation for several future extensions. Incorporating additional modalities—such as multiple morphological stains or the combined gene and protein expression data now available on platforms like Xenium [50]—could further improve segmentation accuracy. Extending to three-dimensional spatial transcriptomics [51, 5], where current 5 *µ*m tissue sections often capture incomplete cells, would enable volumetric cell reconstruction by redefining the high-resolution representations in three-dimensional coordinates. Optimizing gene selection to prioritize spatially representative genes while avoiding those prone to high transcript diffusion [52] could also enhance single-cell information recovery on sequencing-based platforms.

In summary, CellART serves as a practical and versatile tool for extracting single-cell information from high-resolution ST data, distinguished by its robustness, efficiency, and scalability. By transforming raw ST datasets into biologically meaningful cell-level information, CellART provides a critical foundation for downstream analyses, enabling applications such as niche identification, tumor microenvironment characterization, and cell-cell interaction studies. Its unified framework bridges the gap between high-resolution spatial data and cellular-level insights, establishing itself as an essential method for advancing spatial biology across diverse platforms and biological contexts.

## Methods

CellART is a unified framework for extracting single-cell information from high-resolution ST data. The primary objectives are to accurately delineate boundaries for individual cells and further annotate their cell types. Although cell segmentation and cell type annotation may appear distinct, they are inherently interconnected, with each task reinforcing the other. Understanding cell types aids in accurately assigning transcripts, while effective segmentation provides critical spatial information to infer cell types. The integrative application of deep neural networks and probabilistic modeling enables CellART to effectively harness multimodal data, utilizing information from spatial transcriptomics, staining images, and scRNA-seq references to simultaneously achieve accurate cell segmentation and precise cell type annotation, thereby providing a solid foundation for downstream analyses. Moreover, CellART is compatible with a wide range of high-resolution ST platforms, such as Xenium, VisiumHD, Stereo-seq, and MERFISH, making it a versatile and powerful tool in the spatial transcriptomics research.

At a high level, CellART works in three steps. First, it learns a compact latent representation for every subcellular spot by training a feature pyramid network (FPN) whose output is linked, through a Poisson likelihood, to known cell type expression profiles from a scRNA-seq reference. This probabilistic coupling ensures that spots belonging to the same cell type acquire similar representations, even when individual spot measurements are sparse or noisy. Second, CellART uses these biologically informed representations—together with a geometric prior based on proximity to detected nuclei—to classify each spot as belonging to a specific cell, thereby producing smooth cell boundaries. Third, the improved cell boundaries are used to aggregate spot-level information into whole-cell expression profiles, and the probabilistic model is further refined to produce final cell type annotations. The mathematical details of each step are provided below.

### The model of CellART

We denote the observed gene expression count data as **Y** = [*Y*_*i,j,g*_] ∈ ℝ^*H×W ×G*^, where *H* and *W* represent the height and the width of the spatial data, respectively, and *G* corresponds to the number of genes. Each spot in the dataset is indexed by the coordinates (*i, j*), where 1 ≤ *i* ≤ *H* and 1 ≤ *j* ≤ *W* . Each expression matrix is accompanied by a corresponding H&E staining image with dimensions *H* and *W* . To better present our core idea, we assume that the nuclei have been accurately segmented from the staining image using tools such as StarDist or Cellpose. We denote the nuclei segmentation mask as 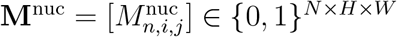, where *N* represents the number of cells. In this context, 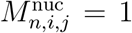 indicates that the spot (*i, j*) belongs to the nucleus of cell *n*, while 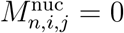 indicates otherwise. Furthermore, a reference matrix ***µ*** = [*µ*_*c,g*_] ∈ ℝ^*C×G*^ that includes cell-type-specific signatures can be constructed from an annotated reference scRNA-seq dataset, where the index *c* = 1, 2, …, *C* denotes the cell type, and each row ***µ***_*c*_ ∈ ℝ^*G*^ represents the mean expression profile of a specific cell type. The reference matrix is normalized such that ∑_*g*_ *µ*_*c,g*_ = 1.

To couple cell segmentation with annotation, we introduce for each spot (*i, j*) a latent representation **Z**_*i,j*_ ∈ ℝ^*D*^ with *D < G*. This shared latent space serves three purposes. First, it reduces sparse high-dimensional gene expression to a compact representation in which similarity between spots (*i, j*) and (*i*^*′*^, *j*^*′*^) is reflected by the proximity of **Z**_*i,j*_ and **Z**_*i*_*′*_,*j*_*′* . Second, it facilitates cell segmentation by grouping spatially adjacent spots with similar latent features, which delineates cell boundaries. Third, after assigning spots to cells, the latent representations of all spots belonging to cell *n* are aggregated to form a cell-level representation **z**_*n*_, which is mapped to a cell type probability vector ***β***_*n*_ = (*β*_*n*,1_, …, *β*_*n,C*_), where *β*_*n,c*_ denotes the probability of cell *n* belonging to type *c*. Cells with similar latent representations tend to have similar cell type probability vectors, allowing for reliable inference of cell types.

An FPN [53] with a ResNet [54] backbone is used to extract high-resolution latent representations **Z** = [*Z*_*i,j,d*_] ∈ ℝ^*H×W ×D*^ from the normalized and log-transformed expression data **X** = [*X*] ∈ ℝ^*H×W ×G*^, where 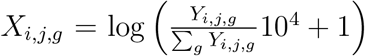. In our framework, the FPN is expected to effectively capture both local and global transcriptomic landscapes within the ST data. The spot-level embeddings are obtained through the FPN as 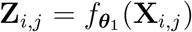, where ***θ***_1_ denotes the network parameters. In the following, we provide a detailed description of the three-step framework designed to extract single-cell-level information from high-resolution ST data, comprising 1) high-resolution representation learning, 2) cell segmentation, and 3) cell type annotation. These steps are illustrated in Figure 1a.

#### Step 1: High-resolution representation learning

To fully leverage subcellular measurements, it is critical to learn the FPN 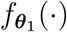 to obtain high-resolution representation **Z**_*i,j*_ for all spots (Fig. 1a: Step 1). The structure of the FPN is illustrated in Supplementary Fig. S34. A naive approach would involve using an autoencoder to reconstruct spot-level gene expression **X**_*i,j*_ from **Z**_*i,j*_. However, this approach is ineffective due to the low transcriptional abundance in individual spots. Instead, we aggregate **Z**_*i,j*_ into a cell-level representation **z**_*n*_ based on the nuclei segmentation. We then integrate the scRNA-seq reference data with **z**_*n*_ to infer the cell type probability vector ***β***_*n*_ using a probabilistic model. This approach allows the reference information to enhance the learning of the FPN, generating accurate representations at both the spot and cell levels, namely **Z**_*i,j*_ and **z**_*n*_.

Specifically, we obtain the cell-level representation as 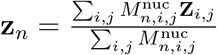 for *n* = 1, 2, …, *N* . Based on the principle that cells with similar representations tend to share the same cell identity, we model the relationship between the cell representation **z**_*n*_ and the cell type probability vector ***β***_*n*_ through a neural network 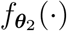, where 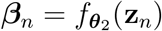. The network is implemented as a multilayer perceptron with a softmax function to enforce 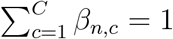 and ***θ***_2_ denotes the network parameters. We denote the observed nucleus gene expression profile for cell *n* as 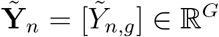, which is obtained by aggregating the raw counts **Y** according to the nuclei segmentation mask **M**^nuc^ as 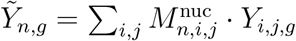.

Next, we introduce a probabilistic model to relate **z**_*n*_ with 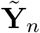, where the reference signature matrix ***µ*** is incorporated. To account for platform discrepancies between ST and scRNA-seq data, we further introduce two effects, denoted as ***α*** = [*α*_*n*_] ∈ ℝ^*N*^ and ***γ*** = [*γ*_*g*_] ∈ ℝ^*G*^. *α*_*n*_ models the variations in total transcript counts across different cells, while *γ*_*g*_ accounts for expression differences of gene *g* between the ST and scRNA-seq platforms. We use a Poisson-based probabilistic model to describe the transcript count 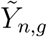 for gene *g* in the nucleus of cell *n* as follows:

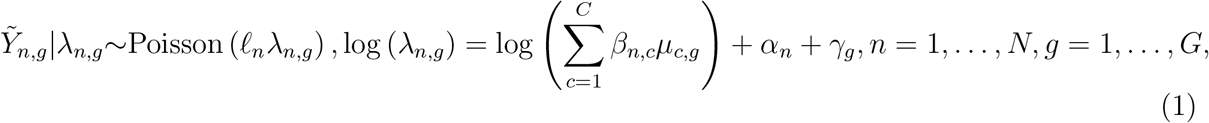

where 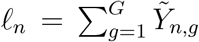 is the total transcript count in the corresponding nucleus. We use the Poisson distribution because it is well accepted as a measurement model for scRNA-seq data and ST data. We have evaluated alternative likelihood formulations, including the Negative Binomial, Zero-Inflated Poisson, Zero-Inflated Negative Binomial, and Tweedie distributions, and found that all alternatives produced virtually identical results in both cell type annotation and segmentation (Supplementary Fig. S43), confirming the adequacy of the Poisson specification for current high-resolution ST platforms.

As *α*_*n*_ models the cell-level variation, we assume that *α*_*n*_ can be extracted from the latent representation **z**_*n*_ through a network parameterized by ***θ***_3_, such that 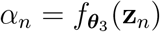. The gene-specific effect *γ*_*g*_ is modeled as a learnable parameter. To ensure that the learned representations retain sufficient information for accurate reconstruction, we consider a decoder network 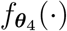 to reconstruct the normalized expression of each cell **X**_*n*_ from the cell representation **z**_*n*_. Therefore, the optimization problem is given as follows:

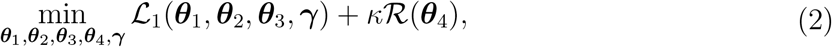

where 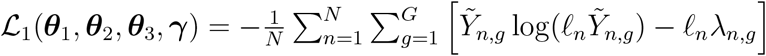 corresponds to the negative Poisson likelihood, 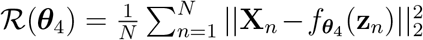 s the regularizer, and *κ* is a regularization parameter fixed to 0.1 throughout this study. We employ stochastic gradient descent to update the model parameters.

#### Step 2: Cell segmentation

With the high-resolution representation **Z**_*i,j*_ obtained in Step 1, we design a segmentation network to classify each spot into its corresponding cell (Fig. 1, Step 2). To obtain the cell probability map, the high-resolution representation is first reduced to a single channel while preserving the spatial dimensions (*i, j*). Since **Z**_*i,j*_ has already captured both local and global features of the ST data through the FPN, the dimensionality reduction is performed using 1*×*1 convolutional layers [55], which maintain computational efficiency and prevent entanglement of the encoded spot-level information.

To better determine which cell each subcellular spot should belong to, CellART introduces a Nucleus Influence Field (NIF) for each potential cell, based on the nuclei segmentation masks **M**^*nuc*^. The NIF is constructed under the assumption that regions closer to the nucleus are more likely to belong to the corresponding cell [56]. Specifically, for a given cell *n*, the NIF is defined as:

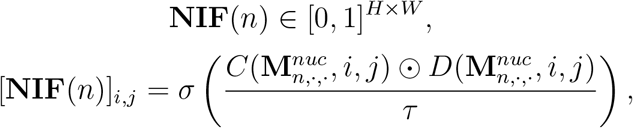

where ⊙ denotes element-wise multiplication. In this formulation, 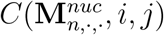 is assigned a value of 1 if the spot (*i, j*) lies within the nucleus of cell *n*, and -1 otherwise. The term 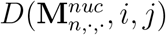 represents the Euclidean distance from spot (*i, j*) to the nearest spot within the nucleus of cell *n*. The parameter *τ* controls the decay rate of the influence field and we set *τ* = 5 throughout our study, which results in reasonable cell sizes across platforms. The sigmoid function *σ*(·) ensures that NIF values are bounded within the range [0,1]. An illustration of the NIF is provided in Supplementary Fig. S33. The single-channel output of the 1 *×* 1 convolutional layer is then multiplied element-wise with **NIF**(*n*), allowing the segmentation process to focus on cell *n*. The resulting output is further normalized to the range [0,1] to produce the final predicted probability map. This operation incorporates positional information encoded in the NIF, effectively associating subcellular spots with their corresponding nuclei while preserving biologically accurate spatial relationships. This mechanism ensures continuity in cell boundaries, maintains appropriate cell sizes, and eliminates the need for labor-intensive post-processing steps commonly required by alternative methods.

Specifically, the segmentation network takes **Z**_*i,j*_ and the NIF for cell *n* as inputs and outputs 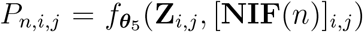, where *P*_*n,i,j*_ denotes the probability that spot (*i, j*) belongs to cell *n*, and ***θ***_5_ represents the parameter set of the deep learning network, which in this case consists solely of 1*×*1 convolutional layers. The architecture of the network is illustrated in Supplementary Fig. S35.

A major challenge in this step is the absence of ground truth labels for each spot, making it difficult to train the segmentation network. To tackle this, we leverage information from the nuclei segmentation mask to generate positive and negative labels. We use spots located within the nuclei as positive samples for each cell, i.e., 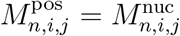. For negative samples, we first dilate each nucleus using a circular kernel with a radius of 10 *µ*m (approximately the cell radius) to create a dilated mask 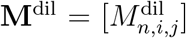. Here, 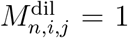 if spot (*i, j*) is within the dilated nucleus of cell *n*, and 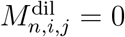 otherwise. Negative samples are defined as spots outside any of the dilated nuclei, such that 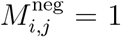 if 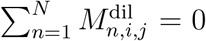, and 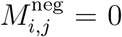 otherwise. This approach ensures that negative samples are sufficiently distant from the nuclei and are likely to be part of the background, where no cell is present.

With high-resolution representations and their corresponding labels, we can train the segmentation network with high accuracy. The rationale is that the spot-level representations from Step 1 are designed to infer cell type information, inherently capturing the spatial distribution of marker genes and the intensity of gene expression. Spots with the same labels are expected to exhibit similar representations. To obtain cell segmentation, we consider the weighted cross entropy loss function to train 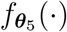:

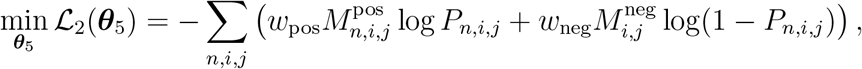

where 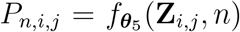, and *w*_pos_ and *w*_neg_ are weights for positive and negative samples, respectively. We set *w*_pos_ = 5 and *w*_neg_ = 1 to address the class imbalance in this study.

After training, we obtain 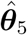 and 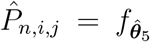 (**Z**_*i,j*_, [**NIF**(*n*)]_*i,j*_). A spot (*i, j*) is classified as background if its probabilities of belonging to all cells are below a threshold *δ* (i.e., 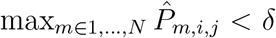), where *δ* = 0.5 in our experiments. The final cell segmentation mask 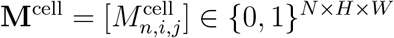 is obtained by assigning each spot to the cell with the highest probability:

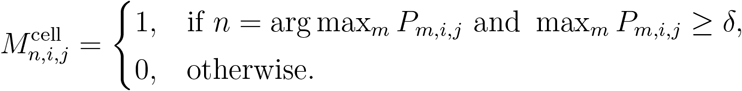

CellART’s boundary determination adapts to the local transcriptional landscape (Fig. 4**e,f** ; Supplementary Fig. S44): when neighboring cells are transcriptionally distinct, boundaries are driven by expression similarity; when neighboring cells are similar in their transcriptional profiles, boundaries are governed by the Nuclei Influence Field geometric prior, producing a Voronoi-like tessellation that ensures each cell retains its complete nucleus and maintains biologically plausible size.

#### Step 3: Cell type annotation

After obtaining the cell segmentation mask **M**^cell^, we update the cell-level representations and expression profiles as 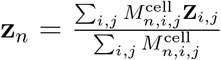 and 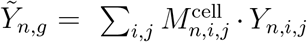. Then, we utilize the same model described in Eq. (1) for cell type annotation (Fig. 1a: Step 3), with all parameters fine-tuned during this step. The key difference from Step 1 is that we now have transcriptomic information for the entire cell rather than just the nucleus. This further reduces measurement noise and improves accuracy in cell type annotation. Finally, we have the inferred cell type probability vector ***β***_*n*_.

### Patchification strategy for large-scale high-resolution ST data

To handle the large-scale nature of high-resolution ST data, CellART employs a patchification strategy (Supplementary Fig. S36). Typically, the spatial dimensions of high-resolution ST data can reach tens of thousands of pixels, making it computationally intensive to process the entire dataset at once. To address this, CellART divides the large ST dataset into smaller, manageable patches. Specifically, the observed gene expression count data **Y** is partitioned into *K* non-overlapping patches of size *h × w*, where *h* and *w* are chosen based on the computational resources available (default values are *h* = *w* = 400). If *H* or *W* is not divisible by *h* or *w*, additional patches are created along the right and bottom edges of the tissue to ensure full coverage. Each patch is indexed by *k* ∈ *{*1, 2, …, *K}*, and the corresponding gene expression data for patch *k* is denoted as 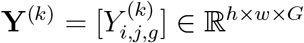. All the network parameters are shared across different patches to ensure consistency in representation learning and segmentation, except that the gene-specific effect ***γ*** is designed to be patch-specific to account for potential batch effects across different patches. Each patch is processed independently through the three-step CellART framework and served like a batch in deep learning (calculate loss and optimize per patch). Note that in the training process, the cells located at the edges of patches may be incomplete, so they are excluded from the loss function calculation to avoid introducing noise. In the prediction phase, to handle cells that span multiple patches, CellART incorporates a shifted patch strategy. This involves creating additional patches that are shifted either horizontally or vertically by half the patch size (i.e., shifted by *h/*2 or *w/*2). By processing these shifted patches, CellART can capture cells that cross the boundaries of the original patches. The final segmentation and cell type annotation results are obtained by merging the outputs from both the original and shifted patches, ensuring comprehensive coverage of all cells in the dataset.

### The reference-free mode of CellART using non-negative matrix factorization

While CellART primarily utilizes scRNA-seq reference data for cell type annotation, it also offers a reference-free mode that leverages non-negative matrix factorization (NMF) to identify distinct cell populations directly from high-resolution ST data (Supplementary Fig. S37). This mode is particularly useful when suitable reference datasets are unavailable or when exploring novel cell types.

In the reference-free mode, CellART replaces the signature matrix *µ* in Eq. (1) with a learnable non-negative matrix **W** = [*W*_*c,g*_] ∈ ℝ^*C×G*^), where each row **W**_*c*_ represents the gene expression profile of cell population *c*. The trainable parameters **W** are optimized alongside other parameters in CellART. The number of cell populations *C* can be specified based on prior knowledge or initially set to a relatively large value, which can then be refined by merging populations based on similarities in their expression profiles or spatial distributions. After training, each cell *n* is assigned to the cell population with the highest probability in its cell type probability vector ***β***_*n*_. This approach enables CellART to uncover underlying cellular heterogeneity in the ST data without relying on external references. To determine the potential cell types represented by each population, we present a workflow that first identifies marker genes for each population and then employs large language models (LLMs) to assist in cell type identification based on these markers. An example using the Xenium human skin cancer dataset is provided in Supplementary Fig. S38.

### Preprocessing

To obtain the input data for CellART (i.e., the observed gene expression count matrix **Y** and the nuclei segmentation mask **M**^nuc^), we need to preprocess the raw high-resolution ST data and segment the nuclei from the staining images. For imaging-based ST platforms such as Xenium and MERFISH, which directly provide the spatial locations of transcripts and therefore do not involve a predefined spot size, we first aggregate the raw transcript coordinates into small, fixed-size spots (1 *µ*m *×* 1 *µ*m) to construct the observed gene expression count matrix **Y**. Next, since the raw dataset typically includes a nuclei segmentation mask as part of its workflow, we directly use it as **M**^nuc^. For sequencing-based ST platforms, the spot sizes vary (e.g., 2 *µ*m for VisiumHD and 0.5 *µ*m for Stereo-seq), and the transcripts are relatively sparse. Therefore, we set the resolution to 2 *µ*m for both VisiumHD and Stereo-seq datasets to balance transcript density and spatial resolution. For VisiumHD datasets, the raw 2 *µ*m spots are directly used as **Y**, and StarDist is applied to segment the nuclei from the H&E staining images. For Stereo-seq datasets, the 0.5 *µ*m spots are binned into 2 *µ*m spots to construct **Y** and Cellpose is used to segment the nuclei from the DAPI staining images. All genes shared between the imaging-based ST data and the scRNA-seq reference are directly used as input features, whereas in sequencing-based ST platforms, the input genes are selected as the intersection between the ST data and the highly variable genes (HVGs) identified from the scRNA-seq reference using the Scanpy package (2,000 HVGs by default). This gene selection strategy for sequencing-based platforms balances transcriptomic coverage with computational efficiency: HVGs from the scRNA-seq reference capture the genes most informative for distinguishing cell types, while the intersection with the ST gene panel ensures that only measurable genes are included. The default of 2,000 HVGs provides sufficient discriminative power for cell type annotation across the tissue types tested in this study, though this number can be adjusted by the user based on the complexity of the tissue and the depth of the scRNA-seq reference.

Before CellART execution, an optional nuclei segmentation quality control and preprocessing pipeline can be applied to the nuclei mask **M**^nuc^. This pipeline includes: (1) filtering of artifact nuclei with extremely small area or negligible UMI counts, (2) an optional multi-nuclei merge step that identifies spatially proximate nuclei with highly correlated gene expression profiles and consolidates them into single cell entities. Details and validation are provided in the Supplementary (Supplementary Figs. S40 and S41).

All processed ST and scRNA-seq reference datasets used in this study are provided in the CellART GitHub repository.

### Data description

#### Xenium 2.0 human lung dataset

The Xenium 2.0 human lung dataset is obtained from the 10x Genomics website (https://www.10xgenomics.com/cn/datasets/preview-data-ffpe-human-lung-cancer-with-xenium-multimodal-cell-segmentation-1-standard). The dataset contains 377 genes measured in a human lung tissue. The area of the tissue is approximately 30 *mm*^2^ with 162,254 cells. This dataset also provides multimodal cell segmentation results with Xenium Onboard Analysis version 2.0.0 from multiple morphological staining images (including the nuclei staining and the cell membrane staining), which is used as the benchmark for comparison.

#### Xenium human lung dataset (Post-Xenium In Situ Applications)

The Xenium human lung dataset is obtained from the 10x Genomics website (https://www.10xgenomics.com/cn/datasets/xenium-human-lung-cancer-post-xenium-technote). This dataset is generated by 10x Genomics in a technical note to demonstrate that the tissue sections processed using the Xenium In Situ workflow remain largely intact post-run and can be used for additional applications. So a paired post-Xenium VisiumHD dataset of the same tissue section is also provided. The Xenium dataset contains 289 genes measured in a human lung tissue. The area of the tissue is approximately 72 *mm*^2^. The nuclei segmentation mask is provided in the raw dataset, resulting in 278,659 segmented nuclei.

#### Xenium mouse brain dataset

The Xenium mouse brain dataset is obtained from the 10x Genomics website (https://www.10xgenomics.com/datasets/fresh-frozen-mouse-brain-replicates-1-standard). The dataset contains 252 genes measured in a mouse brain tissue. The area of the tissue is approximately 60 *mm*^2^ with 162,033 nuclei.

#### Xenium human breast cancer dataset

The Xenium human breast cancer dataset is obtained from the 10x Genomics website (https://www.10xgenomics.com/products/xenium-in-situ/preview-dataset-human-breast). This dataset contains two replicates of human breast cancer tissue sections, each measuring approximately 40 *mm*^2^ with 313 genes measured. The nuclei segmentation mask is provided in the raw dataset, resulting in 167,780 segmented nuclei in the first replicate and 118,75 in the second replicate. Both replicates are analyzed in this study.

#### Xenium colorectal cancer dataset

The Xenium colorectal cancer dataset is obtained from the 10x Genomics website (https://www.10xgenomics.com/platforms/visium/product-family/dataset-human-crc), with name “Xenium In Situ, Sample P2 CRC”. The original complete datasets include three colorectal cancer tissue sections from different patients profiled by both Xenium and VisiumHD platforms. In this study, we only analyze one tissue section (Sample P2). The Xenium dataset of Sample P2 contains 422 genes measured in a colorectal cancer tissue section with an area of approximately 42 *mm*^2^. The nuclei segmentation mask is provided in the raw dataset, resulting in 340,837 segmented nuclei.

#### Xenium human skin dataset

The Xenium human skin dataset is obtained from the 10x Genomics website (https://www.10xgenomics.com/cn/datasets/human-skin-preview-data-xenium-human-skin-gene-expression-panel-add-on-1-standard). This dataset contains a human skin cancer tissue section with an area of approximately 13 *mm*^2^ and 382 genes measured. The nuclei segmentation mask is provided in the raw dataset, resulting in 87,499 segmented nuclei.

#### VisiumHD mouse brain dataset

The VisiumHD mouse brain dataset is obtained from the 10x Genomics website (https://www.10xgenomics.com/cn/datasets/visium-hd-cytassist-gene-expression-libraries-of-mouse-brain-he). This dataset contains a mouse brain tissue section with 6,296,688 2 *µ*m spots and 18,991 genes measured. We use StarDist to segment the nuclei from the matching H&E staining image, resulting in 61,851 segmented nuclei.

#### VisiumHD colorectal cancer dataset

The VisiumHD colorectal cancer dataset is obtained from the 10x Genomics website (https://www.10xgenomics.com/cn/datasets/visium-hd-cytassist-gene-expression-libraries-of-human-colorectal-cancer-tissue), with name “Visium HD, Sample P2 CRC”. This dataset is originally part of a complete dataset that includes three colorectal cancer tissue sections from different patients profiled by both Xenium and VisiumHD platforms. The VisiumHD dataset of Sample P2 contains 8,734,608 2 *µ*m spots and 18,072 genes measured. We use StarDist to segment the nuclei from the matching H&E staining image, resulting in 256,238 segmented nuclei.

#### MERFISH mouse brain dataset

The MERFISH mouse brain dataset is obtained from the Vizgen website (https://info.vizgen.com/mouse-brain-map), with sample name “S2R1”. This dataset contains a mouse brain tissue section with an area of approximately 20 *mm*^2^ and 307 genes measured. The dataset includes 1,823,584 spots with spatial coordinates and gene expression counts. The nuclei segmentation mask is provided in the raw dataset, resulting in 84,566 segmented nuclei.

#### Stereo-seq mouse brain dataset

The Stereo-seq mouse brain dataset is obtained from the BGI website (https://en.stomics.tech/col1241/index.html). The dataset contains a mouse brain tissue section sequencing with a chip of size 1 cm *×* 1 cm and 27,570 genes measured. After binning the raw 0.5 *µ*m spots into 2 *µ*m spots, we obtain 34,574,400 spots. We use Cellpose to segment the nuclei from the registered DAPI staining image, resulting in 79,307 segmented nuclei.

#### scRNA-seq reference datasets

All the scRNA-seq reference datasets used in this study are publicly available. For the human lung tissue, we use lung atlas data from the CZI CELLXGENE data portal: https://cellxgene.cziscience.com/collections/6f6d381a-7701-4781-935c-db10d30de293/. For the human breast cancer tissue, we use the breast cancer scRNA-seq data of the original study, which is available at the 10x Genomics website: https://www.10xgenomics.com/products/xenium-in-situ/preview-dataset-human-breast/.

For the mouse brain tissue, we use a mouse scRNA-seq dataset from the Allen Brain Atlas: https://portal.brain-map.org/atlases-and-data/rnaseq/mouse-whole-cortex-and-hippocampus-smart-seq. For the human colorectal cancer tissue, we use the colorectal cancer scRNA-seq data from the original study, which is available at the 10x Genomics website: https://www.10xgenomics.com/platforms/visium/product-family/dataset-human-crc.

### Benchmarking analysis

We compare CellART with several baseline methods for cell segmentation and cell type annotation on the high-resolution ST data. For cell segmentation, we compare CellART with Cellpose, Stardist, Bin2cell, BIDCell, ProSeg and Baysor. For cell type annotation, we compare CellART with scANVI, Tangram and RCTD.

#### CellART

Most of the hyperparameters of CellART are set to the same default values. The dimension of the high-resolution representation is set to *D* = 128. The patch size is set to 400 *×* 400. The learning rate is set to 1e-3. For cell segmentation, the nucleus influence field decay parameter is set to *τ* = 5 and the dilation kernel radius is set to 10 *µ*m. For Xenium and MERFISH datasets, we set the training epochs to 50 for high-resolution representation learning (Step 1), 10 for segmentation (Step 2), and 150 for cell type annotation (Step 3) by default. For VisiumHD and Stereo-seq datasets, we set the training epochs to 200 for high-resolution representation learning (Step 1), 15 for segmentation (Step 2), and 300 for cell type annotation (Step 3) by default since there are more genes included. The number of training epochs for each step may vary across different datasets; the default setting can be adjusted based on the convergence of the loss function and the visualization of the annotation results in practice.

#### Cellpose

We use Cellpose version 3.1.1.1 for nuclei segmentation from DAPI staining images. The pre-trained model “cyto” is used with the default parameters as we found the “nuclei” model to undersegment or omit a considerable number of nuclei in DAPI images. The diameter parameter is automatically estimated by Cellpose.

#### Stardist

We use Stardist version 0.9.1 for nuclei segmentation from H&E staining image. We use the pre-trained model “2D versatile he” with the default parameters.

#### Bin2Cell

We use Bin2Cell version 0.3.3 for cell segmentation on VisiumHD. The microns per pixel (mpp) parameter is set to 0.5 for the H&E staining image in VisiumHD datasets. StarDist is used to segment nuclei from the H&E staining images as part of the Bin2Cell workflow with same settings as above. Other parameters are set to default following the tutorial of Bin2Cell.

#### BIDCell

We use BIDCell version 1.0.3 for cell segmentation from the high-resolution ST data. Since BIDCell needs extensive external data like predefined positive and negative markers for each cell type, cell type elongation and nuclei annotation, which are not always available in practice, we only compare the segmentation performance of BIDCell with CellART in Xenium breast cancer dataset following the tutorial of BIDCell with default parameters.

#### ProSeg

We use ProSeg version 3.0.10 for cell segmentation from the high-resolution ST data. We follow the tutorial of ProSeg with default parameters to run ProSeg on Xenium datasets. Since the cell boundaries are partially overlapped in ProSeg results, we apply a combination of buffering and simplification to clean and optimize boundaries, ensuring smoother edges and reduced complexity before visualization.

#### Baysor

We use Baysor version 0.7.0 for cell segmentation from the high-resolution ST data. We set the scale parameter to 5 and min molecules per cell to 15. Other parameters are set to their default values following the tutorial of Baysor.

#### scANVI

We use scANVI implemented in the scvi-tools package version 1.1.2 for cell type annotation from the high-resolution ST data. We set n-layers to 2, n-latent to 30, max-epochs to 20, and n-samples-per-label to 100 for training scANVI following the tutorial of scvi-tools.

#### Tangram

We use Tangram version 1.0.4 for cell type annotation from the high-resolution ST data. We keep the default parameters to run Tangram on all datasets.

#### RCTD

We use spacexr package for RCTD method for cell type deconvolution from the high-resolution ST data. We use doublet mode, which assigns one to two cell types per spot. All other parameters are set to their default values, following the tutorial of spacexr.

### Software for downstream analysis

We use several software packages for downstream analysis and visualization.

#### SpatialData

SpatialData is a data framework that comprises a FAIR storage format and a collection of Python libraries for performant access, alignment, and processing of uni- and multi-modal spatial omics datasets. The output of CellART can be easily converted to SpatialData format for further analysis. We use SpatialData version 0.2.5 for most of the visualizations in this study.

#### Scanpy

Scanpy is a scalable toolkit for analyzing single-cell gene expression data built jointly with anndata. We use Scanpy version 1.10.3 for preprocessing gene expression data including normalization, log-transformation, highly variable genes selection and partial visualizations in this study.

#### Squidpy

Squidpy is a tool for the analysis and visualization of spatial molecular data. We use Squidpy version 1.4.1 for some of the spatial visualizations and finding spatially variable genes in this study.

#### Harmonics

Harmonics is a computational framework for characterizing cell niches in spatial omics data. We use Harmonics version 0.0.4 for identifying cell niches in cancer tissue based on the output of CellART in this study. All the parameters are set to default following the tutorial of Harmonics.

#### scVelo

scVelo is a scalable toolkit for RNA velocity analysis in single cells. We use scVelo version 0.3.3 for RNA velocity analysis in the Xenium breast cancer dataset using the subcellular components output from CellART segmentation in this study. All the parameters are set to default following the tutorial of scVelo.

#### DESeq2

DESeq2 [57] is a statistical package for differential gene expression analysis. We use PyDESeq2 version 0.5.2 for differential gene expression analysis on the VisiumHD colorectal cancer dataset to compare CellART with the traditional spot-level analysis. Since DESeq2 is originally designed for bulk RNA-seq data, we create meta-cells by aggregating the expression profiles of all cells with the same cell type.

#### LIANA

LIANA [58] is a computational framework for inferring cell-cell communications from single-cell transcriptomics data. We use LIANA version 0.3.3 for identifying ligand-receptor pairs of TAMs and tumor cells in the VisiumHD colorectal cancer dataset based on the output of CellART in this study. We use the “rank-aggregate” function to get an aggregate of ligand-receptor scores from multiple methods with minimum expression proportion for the ligands and receptors (expr-prop) set to 0.05 and other parameters set to default, following the tutorial of LIANA.

### Simulation framework for subcellular spatial transcriptomics

To enable rigorous evaluation of cell segmentation and annotation accuracy under controlled conditions with known ground truth, we developed a dedicated simulation framework for subcellular spatial transcriptomics data (Supplementary Fig. S49). Existing simulation tools generate cell-level or spot-level expression data and cannot produce spatially resolved subcellular transcript distributions with known ground-truth cell boundaries, necessitating a purpose-built simulator.

#### Cell layout generation

Nucleus centers are sampled via Poisson disk sampling (minimum inter-nucleus distance 12 pixels) on a 2D spatial field of 1,600 *×* 1,600 pixels. Rectangular empty zones and sparse zones (acceptance rate 0.2) are introduced for realism. Each nucleus is rasterized as an elliptical region (semi-axes 5–6 pixels). Full cell boundaries are generated by Voronoi tessellation from nucleus centers, clipped at a maximum cell radius of 10 pixels. Each cell is assigned a ground-truth cell type from *K* = 6 types.

#### Transcript count sampling

A reference basis matrix **B** ∈ ℝ^*K×G*^ (*G* = 500 genes) is constructed with low housekeeping expression and 15 non-overlapping marker genes per type with elevated expression, row-normalized to sum to 1. For each cell *i* of type *k*_*i*_, transcript counts are drawn from 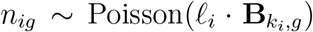, where *ℓ*_*i*_ ∼ LogNormal(log(500), 0.3) is a cell-specific library size.

#### Spatial transcript placement with noise

Each transcript is placed within its parent cell with nuclear enrichment (40% nuclear, 60% cytoplasmic), then displaced by a Gaussian perturbation *N* (0, *σ*^2^**I**) to model boundary diffusion. A fraction *α* of total transcripts is scattered uniformly across the field as ambient background noise, with gene composition drawn from the average expression profile. The resulting gene expression map of shape (*H × W × G*) captures both boundary diffusion and ambient contamination, with *σ* and *α* as tunable parameters.

#### Evaluation

CellART was evaluated in its fully unsupervised NMF mode (no reference expression profiles) with default parameters. Predicted NMF components were aligned to ground-truth cell types using the Hungarian algorithm. Segmentation accuracy was measured by per-cell Intersection over Union (IoU), and annotation accuracy by Hungarian-matched accuracy, Adjusted Rand Index (ARI), and Normalized Mutual Information (NMI). CellART was bench-marked against ProSeg and Baysor on the same simulated data. Systematic parameter sweeps over boundary diffusion (*σ* ∈ *{*0, 2, 4, 6, 8, 10*}* px), ambient noise (*α* ∈ *{*0, 0.05, 0.1, 0.2, 0.3*}*), and transcript density (mean UMI ∈ *{*50, 100, 200, 300, 400, 500*}*) were conducted to evaluate robustness across difficulty regimes. All simulation code and evaluation scripts are available in the code repository.

### Evaluation metrics

We use several metrics to evaluate the performance of cell segmentation and cell type annotation. Since the ground truth cell segmentation masks and the ground truth cell type labels are not always available in real high-resolution ST datasets, we mainly use qualitative evaluation and indirect evaluation based on biological knowledge to assess the performance of different methods in real datasets.

#### F1 score

The F1 score is used to evaluate the accuracy of cell segmentation when the ground truth cell segmentation mask is available. For each cell, we calculate the confusion matrix based on the predicted cell segmentation mask and the ground truth cell segmentation mask. Specifically, the true positive (TP) is defined as the number of spots correctly assigned to the cell, the false positive (FP) is defined as the number of spots incorrectly assigned to the cell, and the false negative (FN) is defined as the number of spots that belong to the cell but are not assigned to it. The precision and recall are calculated as 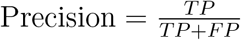 and 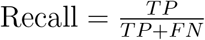. The F1 score is then calculated as the harmonic mean of precision and recall: 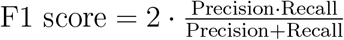. The overall F1 score is obtained by averaging the F1 scores of all cells.

#### Jaccard index

The Jaccard index is used to evaluate the accuracy of cell segmentation when the ground truth cell segmentation mask is available, following the same definition of TP, FP, and FN as above. The Jaccard index is calculated as Jaccard index 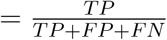. The overall Jaccard index is obtained by averaging the Jaccard indices of all cells.

#### Transcripts cover rate

The transcripts cover rate is used to evaluate the performance of cell segmentation without a ground truth cell segmentation mask. We calculate the proportion of transcripts that are located within the segmented cells. Specifically, we first calculate the total number of transcripts in the tissue, denoted as *T*_total_. Then, we calculate the number of transcripts that are located within the segmented cells, denoted as *T*_in-cell_. The transcripts cover rate is then calculated as Transcripts cover rate 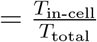. Note that a good cell segmentation method should achieve a high transcripts cover rate while maintaining reasonable cell sizes and high molecule homogeneity within cells.

#### Component correlation

The component correlation is used to compare the molecule heterogeneity within cells between different segmentation methods. For each segmented cell, we calculate the Pearson correlation coefficient between the gene expression profile of the cytoplasm and that of the nucleus. Specifically, for each cell *n*, we denote the gene expression profile of the cytoplasm as 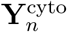 and that of the nucleus as 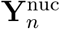. The component correlation for cell *n* is then calculated as Component 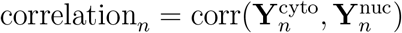. The overall component correlation is obtained by averaging the component correlations of all cells. A good segmentation method should produce high component correlation, indicating that the gene expression profiles of the cytoplasm and nucleus are consistent within each cell.

#### Positive marker purity

The positive marker purity is used to evaluate the accuracy of both cell segmentation and cell type annotation without ground truth cell type labels by leveraging markers defined from the scRNA-seq reference. For each cell type, we first define the positive markers as the top 10 percent highly expressed genes expressed in at least 50 percent of the cells in that cell type in the scRNA-seq reference. Then, for each annotated cell, we calculate the proportion of positive markers that are expressed in the cell as the sensitivity, and use box plots to visualize the distribution of sensitivity across all cells. A good cell type annotation and segmentation method should produce high positive marker purity, indicating that the annotated cells accurately reflect the gene expression profiles of their respective cell types.

#### Correlation with the reference

The correlation with reference is used to evaluate the accuracy of cell type annotation without ground truth cell type labels. For each annotated cell type, we calculate the Pearson correlation coefficient between the average gene expression profile of the annotated cells and that of the corresponding cell type in the scRNA-seq reference. Specifically, for each cell type *c*, we denote the average gene expression profile of the annotated cells as 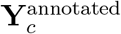 and that of the corresponding cell type in the reference as 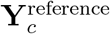. The correlation with reference for cell type *c* is then calculated as Correlation with 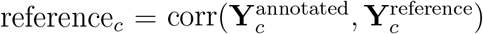. The overall correlation with reference is obtained by averaging the correlations of all cell types. A good cell type annotation method should produce high correlation with the reference, indicating that the annotated cells accurately reflect the gene expression profiles of their respective cell types.

#### Eccentricity

Eccentricity is used to evaluate the shape of segmented cells. For each cell type, we calculate the average eccentricity of all cells and nuclei. Specifically, the eccentricity of an object is defined as the ratio of the distance between the foci of the ellipse that best fits the object and its major axis length. For each cell and nucleus, we first fit an ellipse to its shape and then calculate its eccentricity. A good segmentation method should produce cells with eccentricity similar to that of nuclei, and have diversity in eccentricity across different cell types.

## Code availability

The CellART software is available at https://github.com/YangLabHKUST/CellART.

## Acknowledgements

This work was supported in part by the Innovation and Technology Commission (ITCPD/17-9); Hong Kong Research Grants Council Grants, AoE/E-601/24-N, C6040-24G, 16308120, 16307221, 16307322, 16302823, 16309424, and 16308925; The Hong Kong University of Science and Technology Startup Grants R9405 and Z0428 from the Big Data Institute. The computation tasks for this work were performed using the X-GPU cluster supported by the Research Grants Council Collaborative Research Fund Grant C6021-19EF and HKUST SuperPOD. J.X. was supported by National Natural Science Foundation of China (Grant No. 12401384); Guangdong Natural Science Foundation General Project (Grant No. 2025A1515011603); Sun Yat-sen University Startup Grant (Grant No. 2026 51000 B26833); and Shenzhen Science and Technology Program (Grant No. RCBS20231211090613024).

## Author contributions

C. Y. conceived the study and supervised the project. Y. C. designed, implemented, and validated CellART. J. X. and Y. L. contributed to the experimental design. Y. Z. and Z. W. assisted with the validation of CellART. Z. C., P. J., H. C., and J. W. helped with analyzing the results. Y. C. and C. Y. wrote the manuscript with input from all the authors.

## References

[1] Marx, V. Method of the year: spatially resolved transcriptomics. Nature methods 18, 9–14 (2021).

[2] Eisenstein, M. Seven technologies to watch in 2024. Nature 625, 844–848 (2024).

[3] Oliveira, M. F. d. et al. High-definition spatial transcriptomic profiling of immune cell populations in colorectal cancer. Nature Genetics 1–12 (2025).

[4] Cho, C.-S. et al. Microscopic examination of spatial transcriptome using seq-scope. Cell 184, 3559–3572 (2021).

[5] Schott, M. et al. Open-ST: High-resolution spatial transcriptomics in 3d. Cell 187, 3953–3972 (2024).

[6] Chen, A. et al. Spatiotemporal transcriptomic atlas of mouse organogenesis using dna nanoball-patterned arrays. Cell 185, 1777–1792 (2022).

[7] Janesick, A. et al. High resolution mapping of the tumor microenvironment using integrated single-cell, spatial and in situ analysis. Nature Communications 14, 8353 (2023).

[8] Chen, K. H., Boettiger, A. N., Moffitt, J. R., Wang, S. & Zhuang, X. Spatially resolved, highly multiplexed rna profiling in single cells. Science 348, aaa6090 (2015).

[9] Williams, C. et al. Spatial insights into tumor immune evasion illuminated with 1000-plex rna profiling with cosmx spatial molecular imager. Cancer Res 83, 10 (2023).

[10] Wang, G. et al. Construction of a 3D whole organism spatial atlas by joint modelling of multiple slices with deep neural networks. Nature Machine Intelligence 5, 1200–1213 (2023).

[11] Wan, X. et al. Integrating spatial and single-cell transcriptomics data using deep generative models with SpatialScope. Nature Communications 14, 7848 (2023).

[12] Wang, Z. et al. A unified framework for identification of cell-type-specific spatially variable genes in spatial transcriptomic studies. Proceedings of the National Academy of Sciences 122, e2503952122 (2025).

[13] Zhao, J. et al. Adversarial domain translation networks for integrating large-scale atlas-level single-cell datasets. Nature Computational Science 2, 317–330 (2022).

[14] Cable, D. M. et al. Robust decomposition of cell type mixtures in spatial transcriptomics. Nature Biotechnology 40, 517–526 (2022).

[15] Kleshchevnikov, V. et al. Cell2location maps fine-grained cell types in spatial transcriptomics. Nature biotechnology 40, 661–671 (2022).

[16] Benjamin, K. et al. Multiscale topology classifies cells in subcellular spatial transcriptomics. Nature 630, 943–949 (2024).

[17] Chen, H., Li, D. & Bar-Joseph, Z. SCS: cell segmentation for high-resolution spatial transcriptomics. Nature methods 20, 1237–1243 (2023).

[18] Polański, K. et al. Bin2cell reconstructs cells from high resolution Visium HD data. Bioinformatics 40, btae546 (2024).

[19] Stringer, C., Wang, T., Michaelos, M. & Pachitariu, M. Cellpose: a generalist algorithm for cellular segmentation. Nature methods 18, 100–106 (2021).

[20] Schmidt, U., Weigert, M., Broaddus, C. & Myers, G. Cell detection with star-convex polygons. In International Conference on Medical Image Computing and Computer-Assisted Intervention, 265–273 (Springer, 2018).

[21] Jones, D. C. et al. Cell simulation as cell segmentation. Nature Methods 1–12 (2025).

[22] Petukhov, V. et al. Cell segmentation in imaging-based spatial transcriptomics. Nature biotechnology 40, 345–354 (2022).

[23] Jin, K. et al. Bering: joint cell segmentation and annotation for spatial transcriptomics with transferred graph embeddings. Nature Communications 16, 6618 (2025).

[24] Pang, M., Roy, T. K., Wu, X. & Tan, K. CelloType: a unified model for segmentation and classification of tissue images. Nature methods 22, 348–357 (2025).

[25] Marconato, L. et al. Spatialdata: an open and universal data framework for spatial omics. Nature methods 22, 58–62 (2025).

[26] Wolf, F. A., Angerer, P. & Theis, F. J. Scanpy: large-scale single-cell gene expression data analysis. Genome biology 19, 15 (2018).

[27] Palla, G. et al. Squidpy: a scalable framework for spatial omics analysis. Nature methods 19, 171–178 (2022).

[28] 10x Genomics. Xenium human lung dataset. https://www.10xgenomics.com/cn/datasets/preview-data-ffpe-human-lung-cancer-with-xenium-multimodal-cell-segmentation-1-standard (2024).

[29] 10x Genomics. Visiumhd mouse brain dataset. https://www.10xgenomics.com/cn/datasets/visium-hd-cytassist-gene-expression-libraries-of-mouse-brain-he (2024).

[30] 10x Genomics. Xenium human lung dataset (post-xenium in situ applications). https://www.10xgenomics.com/cn/datasets/xenium-human-lung-cancer-post-xenium-technote (2025).

[31] 10x Genomics. Xenium mouse brain dataset. https://www.10xgenomics.com/datasets/fresh-frozen-mouse-brain-replicates-1-standard (2023).

[32] Vizgen. Merfish mouse brain dataset. https://info.vizgen.com/mouse-brain-map (2024).

[33] BGI. Stereo-seq mouse brain dataset. https://en.stomics.tech/col1241/index.html (2024).

[34] Wang, Q. et al. The Allen mouse brain common coordinate framework: a 3D reference atlas. Cell 181, 936–953 (2020).

[35] Bird, C. M. & Burgess, N. The hippocampus and memory: insights from spatial processing. Nature reviews neuroscience 9, 182–194 (2008).

[36] Zhang, M. et al. Molecularly defined and spatially resolved cell atlas of the whole mouse brain. Nature 624, 343–354 (2023).

[37] Lopez, R., Regier, J., Cole, M. B., Jordan, M. I. & Yosef, N. Deep generative modeling for single-cell transcriptomics. Nature methods 15, 1053–1058 (2018).

[38] Fu, X. et al. Bidcell: Biologically-informed self-supervised learning for segmentation of subcellular spatial transcriptomics data. Nature communications 15, 509 (2024).

[39] Biancalani, T. et al. Deep learning and alignment of spatially resolved single-cell transcriptomes with tangram. Nature methods 18, 1352–1362 (2021).

[40] Hadler-Olsen, E.Winberg, J.-O. & Uhlin-Hansen, L. Matrix metalloproteinases in cancer: their value as diagnostic and prognostic markers and therapeutic targets. Tumor Biology 34, 2041–2051 (2013).

[41] Serrano-Gomez, S. J., Maziveyi, M. & Alahari, S. K. Regulation of epithelial-mesenchymal transition through epigenetic and post-translational modifications. Molecular cancer 15, 18 (2016).

[42] Sidenius, N. & Blasi, F. The urokinase plasminogen activator system in cancer: recent advances and implication for prognosis and therapy. Cancer and Metastasis Reviews 22, 205–222 (2003).

[43] Desgrosellier, J. S. & Cheresh, D. A. Integrins in cancer: biological implications and therapeutic opportunities. Nature Reviews Cancer 10, 9–22 (2010).

[44] Martens, Y. A. et al. Apoe cascade hypothesis in the pathogenesis of alzheimer’s disease and related dementias. Neuron 110, 1304–1317 (2022).

[45] Dodson, S. E. et al. Loss of lr11/sorla enhances early pathology in a mouse model of amyloidosis: evidence for a proximal role in alzheimer’s disease. Journal of Neuroscience 28, 12877–12886 (2008).

[46] Wang, C. et al. Gain of toxic apolipoprotein e4 effects in human ipsc-derived neurons is ameliorated by a small-molecule structure corrector. Nature medicine 24, 647–657 (2018).

[47] Zalocusky, K. A. et al. Neuronal apoe upregulates mhc-i expression to drive selective neurodegeneration in alzheimer’s disease. Nature neuroscience 24, 786–798 (2021).

[48] Pietilä, M. et al. Sorla regulates endosomal trafficking and oncogenic fitness of her2. Nature communications 10, 2340 (2019).

[49] Zheng, Z. et al. Sorl1 stabilizes abcb1 to promote cisplatin resistance in ovarian cancer. Functional & Integrative Genomics 23, 147 (2023).

[50] 10x Genomics. Xenium in situ gene and protein expression data for ffpe human renal cell carcinoma. https://www.10xgenomics.com/cn/datasets/xenium-protein-ffpe-human-renal-carcinoma (2025).

[51] Yapp, C. et al. Highly multiplexed 3d profiling of cell states and immune niches in human tumors. Nature Methods 1–14 (2025).

[52] Ni, Z. et al. Spotclean adjusts for spot swapping in spatial transcriptomics data. Nature Communications 13, 2971 (2022).

[53] Lin, T.-Y. et al. Feature pyramid networks for object detection. In Proceedings of the IEEE conference on computer vision and pattern recognition, 2117–2125 (2017).

[54] He, K., Zhang, X., Ren, S. & Sun, J. Deep residual learning for image recognition. In Proceedings of the IEEE conference on computer vision and pattern recognition, 770–778 (2016).

[55] Lin, M., Chen, Q. & Yan, S. Network in network. arXiv preprint arXiv:1312.4400 (2013).

[56] Chen, Y., Xu, X., Wan, X., Xiao, J. & Yang, C. UCS: a unified approach to cell segmentation for subcellular spatial transcriptomics. bioRxiv 2024–07 (2024).

[57] Love, M. I., Huber, W. & Anders, S. Moderated estimation of fold change and dispersion for rna-seq data with deseq2. Genome biology 15, 550 (2014).

[58] Dimitrov, D. et al. Comparison of methods and resources for cell-cell communication inference from single-cell rna-seq data. Nature communications 13, 3224 (2022).

